# Mutagenesis Sensitivity Mapping of Human Enhancers *In Vivo*

**DOI:** 10.1101/2024.09.06.611737

**Authors:** Michael Kosicki, Boyang Zhang, Anusri Pampari, Jennifer A. Akiyama, Ingrid Plajzer-Frick, Catherine S. Novak, Stella Tran, Yiwen Zhu, Momoe Kato, Riana D. Hunter, Kianna von Maydell, Sarah Barton, Erik Beckman, Anshul Kundaje, Diane E. Dickel, Axel Visel, Len A. Pennacchio

**Affiliations:** Environmental Genomics & System Biology Division, Lawrence Berkeley National Laboratory, One Cyclotron Road, Berkeley, CA 94720, USA; Department of Genetics, Stanford University, Stanford, CA, USA; Department of Computer Science, Stanford University, Stanford, CA, USA; School of Natural Sciences, University of California, Merced, CA 95343, USA; U.S. Department of Energy Joint Genome Institute, One Cyclotron Road, Berkeley, CA 94720, USA; Comparative Biochemistry Program, University of California, Berkeley, CA 94720, USA

## Abstract

Distant-acting enhancers are central to human development. However, our limited understanding of their functional sequence features prevents the interpretation of enhancer mutations in disease. Here, we determined the functional sensitivity to mutagenesis of human developmental enhancers *in vivo*. Focusing on seven enhancers active in the developing brain, heart, limb and face, we created over 1700 transgenic mice for over 260 mutagenized enhancer alleles. Systematic mutation of 12-basepair blocks collectively altered each sequence feature in each enhancer at least once. We show that 69% of all blocks are required for normal *in vivo* activity, with mutations more commonly resulting in loss (60%) than in gain (9%) of function. Using predictive modeling, we annotated critical nucleotides at base-pair resolution. The vast majority of motifs predicted by these machine learning models (88%) coincided with changes to *in vivo* function, and the models showed considerable sensitivity, identifying 59% of all functional blocks. Taken together, our results reveal that human enhancers contain a high density of sequence features required for their normal *in vivo* function and provide a rich resource for further exploration of human enhancer logic.

## Introduction

Distant-acting enhancers are critical for regulating gene expression in a tissue-specific manner during mammalian development. Enhancer sequences function by binding transcription factors (TFs), proteins that influence the transcriptional output of the enhancer’s target gene^1^. Individual TF binding motifs are typically 6-12bp in size^1^ and most mammalian enhancers are hundreds of basepairs long, containing multiple TF binding sites^2–4^. The potential TF binding sites within an enhancer can be predicted from DNA sequence^2^ and TF binding to DNA in a given tissue or cell type can be directly measured using epigenomic methods such as ChIP-seq^5^. However, given our lack of information on all possible TF binding events, their individual functional contributions, and interactions between bound TFs, we cannot currently predict enhancer activity directly from DNA sequence. This lack of knowledge about the functional underpinnings of enhancers precludes us from predicting how genetic variants affect gene expression.

Enhancer reporter assays offer a way to study the functional relevance of individual subregions or basepairs within an enhancer by coupling wild-type or mutated versions of an enhancer to a reporter gene and measuring the resulting expression. Crucially, these assays allow dissection of enhancer function outside of the enhancer’s endogenous genomic context, where interactions with promoters and other enhancers may confound the readout or even completely mask changes in their individual activity due to enhancer redundancy^6,7^. Recently improved mouse transgenic engineering approaches have enabled larger-scale, whole-organism, sensitive, and reproducible assessment of regulatory elements and mutation effects in the context of prenatal *in vivo* development (enSERT)^8,9^. Changes to spatiotemporal enhancer activity patterns observed in these assays are highly informative of the phenotypic impact of studied mutations on complex processes such as limb or brain development^8,10^. While other, complementary methods for enhancer perturbation (including massively parallel reporter assays) exist, they tend to rely on *in vitro* cell culture systems^11^. Transgenic mouse assays are unique in their ability to reveal the impact of sequence changes within enhancers on their complex spatiotemporal *in vivo* activity patterns in embryonic development.

In the present study, we applied these recent advances in mouse reporter assay technology at scale to explore the sequence determinants of human developmental enhancer function *in vivo*. We conducted a complete, systematic mutagenesis mapping of seven human enhancers active during embryonic development and assessed the consequences of mutations for *in vivo* enhancer activity in mouse transgenic assays. We observed a high density of sites required for correct tissue-specific activity within the enhancers studied, as well as different modes of functional interactions between sites within enhancers. We also trained machine learning models based on chromatin accessibility to predict the binding site motifs within these enhancers and validated them using *in vivo* transgenic assays. The models identified sequence motifs which coincided to a high degree with functional sites, offering a method to computationally predict nucleotides within enhancers that are likely to affect their *in vivo* function. Thus, these models are expected to be useful for the interpretation and prioritization of clinically observed variants in enhancers. Taken together, our data reveal a considerable functional complexity of human *in vivo* enhancers and provide a comprehensive resource for model development and validation.

## Results

### Large-Scale Block Mutagenesis of Developmental Enhancers

To study how the sequence features within mammalian enhancers relate to their *in vivo* activity patterns, we selected seven human enhancers that were between 223bp and 431bp long. Each of these enhancers drives strong and highly reproducible activity in transgenic mouse reporter assays at mid-gestation (embryonic day 11.5) in brain (enhancers NEU1-3), heart (enhancers HT1-3), or face and limb (enhancer FL, Figure 1A, Supplementary Table1)^12–18^. We divided each enhancer into consecutive 12bp blocks for mutagenesis, corresponding to the average size of individual transcription factor binding sites, without biasing the design towards predicted binding sites (Figure 1B). In total, the seven enhancers encompassed 167 mutagenesis blocks. For each enhancer, we generated a series of transgenic reporter constructs in which all basepairs within one or several blocks were mutated using a transition mutagenesis scheme designed to eliminate any transcription factor binding sites that may be present with the block (A<>G, C<>T; Supplementary Figure 1; Supplementary Note1; Supplementary Table2).

**Figure 1.**
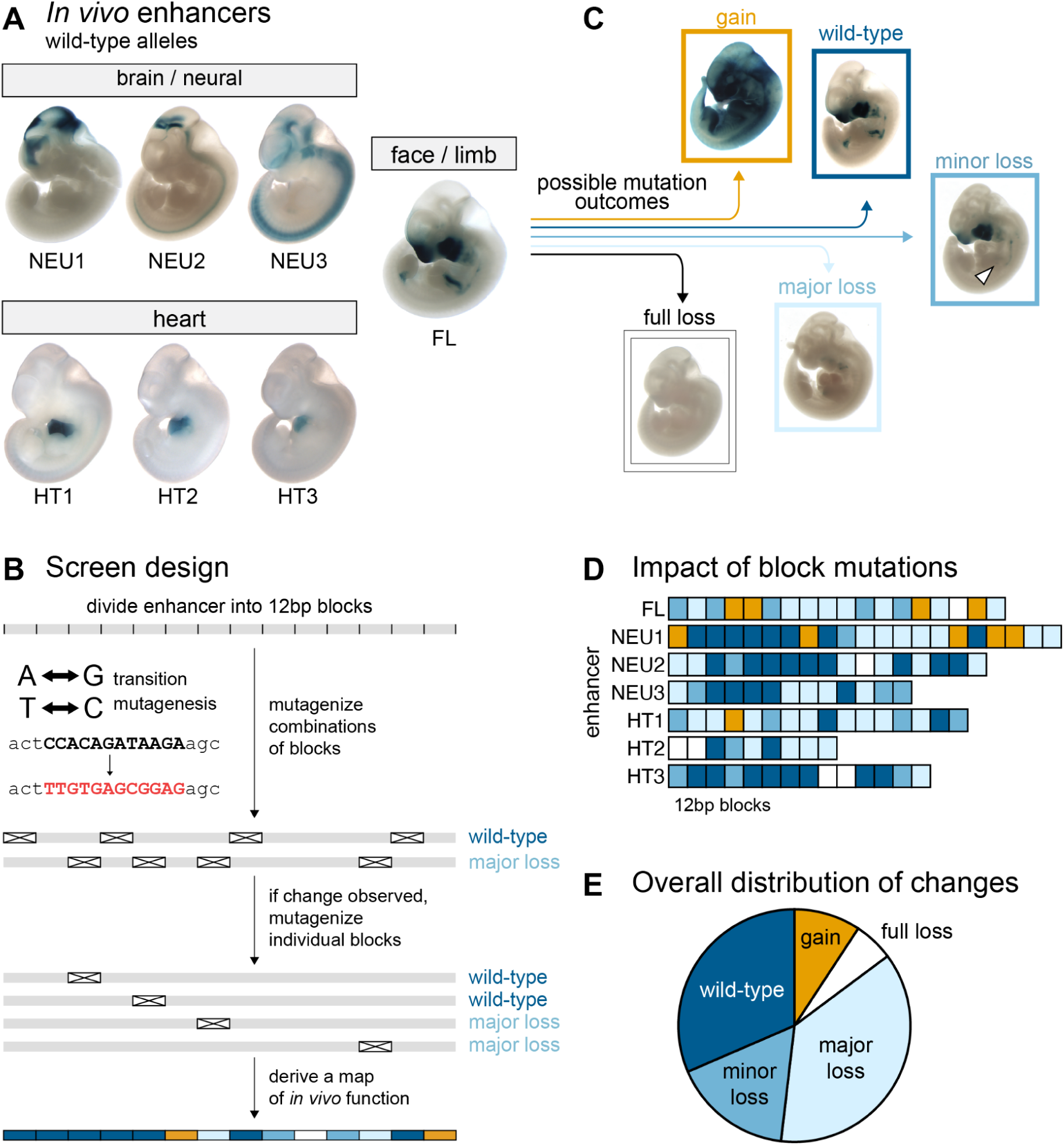
General enhancer properties. (A) Wild-type pattern of seven enhancers mutagenized in this study (see Supplementary Table 1 for details). (B) Initial screen design. (C) Examples of patterns in mutagenized constructs. (D) Functional annotation of 12bp blocks (N=108; see Supplementary Note 2 for adjustments). (E) Distribution of block mutation outcomes (N=108).

To identify subregions of enhancers not required for *in vivo* function, we first produced a series of 103 constructs in which between two and nine 12bp blocks had been mutated simultaneously (Figure 1B). Each mutagenized enhancer was coupled to a minimal promoter and LacZ reporter gene and used to generate transgenic mouse embryos using CRISPR-mediated insertion at a safe harbor locus^8,9^ (enSERT; Methods). We then compared the resulting *in vivo* activity patterns with those of the wildtype allele of each enhancer (Figure 1C). Overall, 33 of the 112 combinatorial constructs, encompassing 69 of the 167 individual blocks, caused no detectable changes in enhancer activity. The absence of changes could theoretically result from combinatorial compensation between loss- and gain-of-function effects. To exclude this possibility, we also tested 21 of these 69 blocks individually in single-block mutation constructs and observed that none of them altered the enhancer activity. Thus, we tentatively classified all 69 blocks as non-critical for *in vivo* enhancer activity. To complete the systematic block mutagenesis survey, we assayed the remaining 98 untested blocks individually, finding an additional 25 non-critical blocks for a total of 94 that appeared dispensable for normal enhancer function. Disruption of the remaining 73 blocks resulted in changes in activity. We also performed additional validation of the transition mutagenesis scheme, which resulted in minor adjustments to functional block annotations (Supplementary Note 2; Supplementary Figure 2).

We observed that the peripheral blocks of many enhancers were often not required for function and therefore we defined the functional core of each enhancer by the two outermost blocks whose mutation caused a change in function (Figure 1D). Across seven enhancers, there was a total of 108 functional core blocks. Mutagenesis of 6% of these 108 blocks led to full loss of function, 37% led to major loss, 17% led to minor loss, 9% led to gain of function and no change was observed when mutagenizing 31% (Figure 1E; Supplementary Table3).

While all seven enhancers contained subregions that caused major changes in activity when mutated, across enhancers we observed notable differences in the proportion of blocks with critical functions and in the types of observed activity changes (Figure 1D). Gain-of-function changes in activity were almost exclusively observed in enhancers FL and NEU1, with 9 of 10 instances located in these two enhancers. This observation suggests that these enhancers contain multiple binding sites for repressive factors that, when mutated, cause de-repression of the enhancer and thereby ectopic activity. Four enhancers (FL, NEU2, HT2, HT3) contained at least one “Achilles’ heel” block that, when mutated, caused a full loss of enhancer function. We also observed substantial differences in the proportion of blocks within an enhancer causing major or full loss of function, ranging from 21% (HT3) to 67% (HT2). Nevertheless, all enhancers contained three or more such blocks.

To investigate whether experimentally observed function agrees with other indicators of DNA function, we examined its relationship with measures of selective constraints in mammalian evolution and in human populations. Blocks that altered *in vivo* enhancer function showed higher evolutionary conservation across mammals than those whose mutation did not cause activity changes (p<0.05, see Supplementary Figure 3A,B,C). Similarly, enhancers with a higher fraction of blocks that caused full/major loss or gain of function showed a lower density of variants across human populations (R^2^=68%, p-value<0.05, Supplementary Figure 3D). These findings support that blocks that contribute to enhancer activity, as observed by mutagenesis screening, contribute to fitness and are therefore subject to selective constraints in evolution and human populations. Taken together, these results show that all tested enhancers have multiple sites critical for their function, dispersed across extended core regions ranging from approximately 110bp to 250bp in length. However, they show substantial differences in their robustness to mutation and in their propensity to gain or lose tissue-specific activities upon mutation.

### Basepair Resolution Prediction of Critical Sites Within Enhancers

The comprehensive *in vivo* dataset of block-mutated enhancers offers a unique opportunity to develop and assess tools for predicting the importance of individual nucleotides for normal *in vivo* enhancer function. We trained a machine learning model (ChromBPNet^19^) to predict genome-wide open chromatin signal from DNA sequence using 29 bulk ATAC-seq, single-cell ATAC-seq (scATAC) and DNase I hypersensitive site sequencing (DHS) human and mouse datasets from embryonic tissues in which the tested enhancers were active (Supplementary Table 4). Next, we used these models to predict the consequences of mutating individual or multiple 12bp blocks in each enhancer and compared the predicted changes in open chromatin signal to the observed differences in enhancer *in vivo* activity. For example, for enhancer FL and using a model derived from e11.5 limb DHS data, mutagenesis of block 12 resulted in a predicted minor reduction (log2 fold change = –0.24) in chromatin openness, which coincided with a minor loss of *in vivo* function in the limbs (Figure 2A). In contrast, mutagenesis of block 16 was predicted to reduce chromatin openness substantially (log2 fold change = –1.03), which coincided with an observed major loss of *in vivo* activity. Comparing all predicted changes in chromatin openness with observed *in vivo* results for enhancer FL revealed a strong correlation (R^2^=0.73, Figure 2B, see Methods for scoring of *in vivo* results). For five of the seven enhancers, we identified models trained on data from relevant tissues with high correlation between predicted mutation effects and *in vivo* results observed for mutant alleles (respective best-fit models: R^2^=0.50-0.79; Methods, Supplementary Figure 4A, Supplementary Table 4, Supplementary Note3). For two of the seven enhancers none of the models from relevant tissues showed good correlation with *in vivo* results and these enhancers were excluded from further analysis (NEU1 and NEU2, see Supplementary Note3 for details).

**Figure 2.**
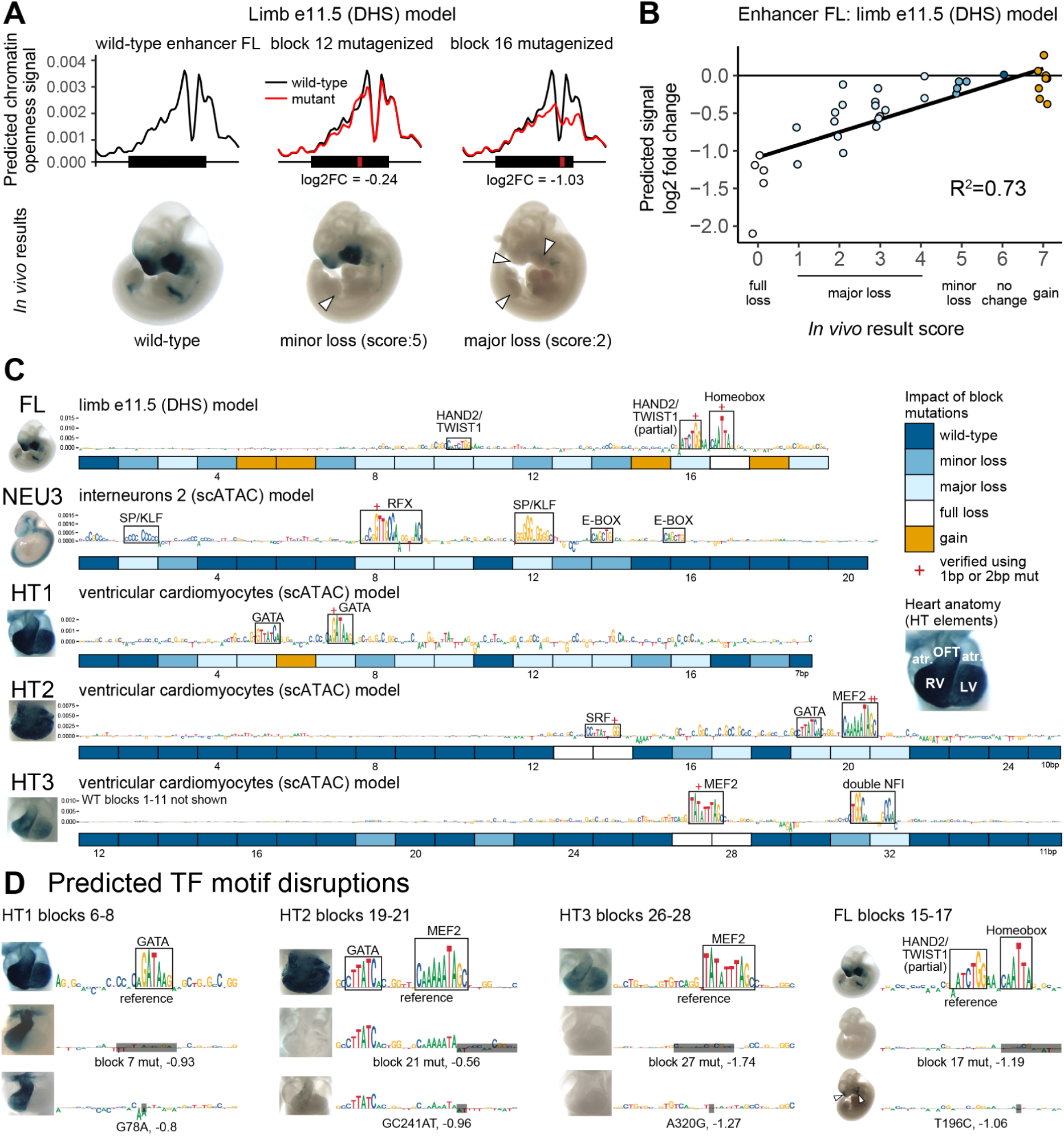
Machine learning model selection and validation. (A) Examples of ChromBPNet model output and *in vivo* results for reference and mutagenized constructs of enhancer FL. White arrowheads indicate partial or full loss of *in vivo* activity. (B) Correlation of model-predicted mutation effects (change in predicted signal between wild-type and mutagenized sequence) and the observed *in vivo* mutagenesis results. Each dot represents a construct with a mutagenized block or a combination of blocks. R^2^=Spearman correlation. (C) Contributions scores for wild-type sequences with per block *in vivo* experiment results in boxes below. Best-fit models depicted. Clusters with high contribution scores boxed in (N=14). OFT = outflow tract, LV = left ventricle, RV = right ventricle, atr. = atrium. (D) Single or double basepair mutations were introduced at clusters with high contribution scores. Also see Supplementary Figure 4B.

For each model, we used DeepLIFT^20^ to predict the contribution of each basepair within the enhancer to the open chromatin signal (Figure 2C). Using only the best-fit model for each enhancer, we observed 15 locally dense clusters of contiguous nucleotides with high contribution scores. In many cases, the observed clusters were reminiscent of binding motifs of TFs expected to be active in the tissues observed *in vivo.* For example, in face and limb enhancer FL, the approach revealed high contribution scores for motifs relevant to craniofacial and limb development, including an isolated HAND2/TWIST1 E-box motif and a pair of a homeobox and a HAND2/TWIST1 motifs resembling a previously described Coordinator motif (Figure 2C)^21–24^. Likewise, in heart enhancers HT1, HT2 and HT3 we observed clusters of high contribution scores that corresponded to binding motifs for GATA, MEF2, and SRF, all of which are involved in cardiac development (Figure 2C)^22,25^. Of 15 clusters with high contribution scores, 14 overlapped blocks that showed loss of activity upon mutagenesis, indicating high positive predictive value (93%). Conversely, of the 53 blocks whose mutation caused a change of *in vivo* function, 19 overlapped clusters with high contribution scores, indicating moderate sensitivity (36%, also see Supplementary Note 4, Methods).

Next, we assessed experimentally if the motifs identified by high contribution scores are indeed the critical functional components of the 12 basepair blocks tested previously by block mutagenesis. We introduced targeted mutations of single or two adjacent nucleotides predicted to disrupt 7 of 15 clusters with high contribution scores. In all cases, we observed a loss-of-function in line with contribution score-based predictions. For example, in enhancer HT1, upon introducing a point mutation (G78A) within a predicted GATA binding motif, we observed a complete loss of *in vivo* activity in the left cardiac ventricle that was indistinguishable from the effect of mutating the entire surrounding 12 basepair block (Figure 2D). Similar effects were observed for all other cases tested (Figure 2D, Supplementary Figure 4B). Together, these results indicate that contribution scores derived from models trained to predict open chromatin signal can identify functional TF motifs within enhancers and predict the impact of their mutation on enhancer activity with considerable accuracy.

### Consideration of Degenerate Motifs and Multi-Tissue Activities Improves Detection Sensitivity

To increase the sensitivity of detecting functionally relevant TF motifs, we hypothesized that motifs with weaker contribution scores may escape detection because they do not stand out as distinct clusters in wildtype sequence. To find such degenerate sites, we performed *in silico* saturation mutagenesis of all five enhancers, generating 5082 variant sequences with 1bp substitution mutation each. Next, we examined the variant sequences for the emergence of new local clusters of nucleotides with high contribution scores, and for changes in overall predicted open chromatin signal across the enhancer. For example, in enhancer HT1, we observed that a single *in silico* point mutation (T111C) resulted in the emergence of a strong, predicted MEF2 motif that is not evident from the reference sequence. The mutation increased the predicted open chromatin signal substantially (log2 fold change = 0.74; Figure 3A, left). Targeted disruption of this MEF2 motif through mutation of a different single basepair (T112C) caused region-specific loss of cardiac *in vivo* activity in a pattern that was identical to the loss of activity observed upon mutating the entire 12bp block in which the mutation resides. A degenerate MEIS-TEAD site with similar *in vivo* impact was observed in another block of enhancer HT1 (Figure 3A, right). Across all enhancers, we identified 6 sites that both featured a novel cluster of high, positive contribution scores and had a predicted open chromatin signal 25% higher than the reference (Supplementary Figure 5A).

**Figure 3.**
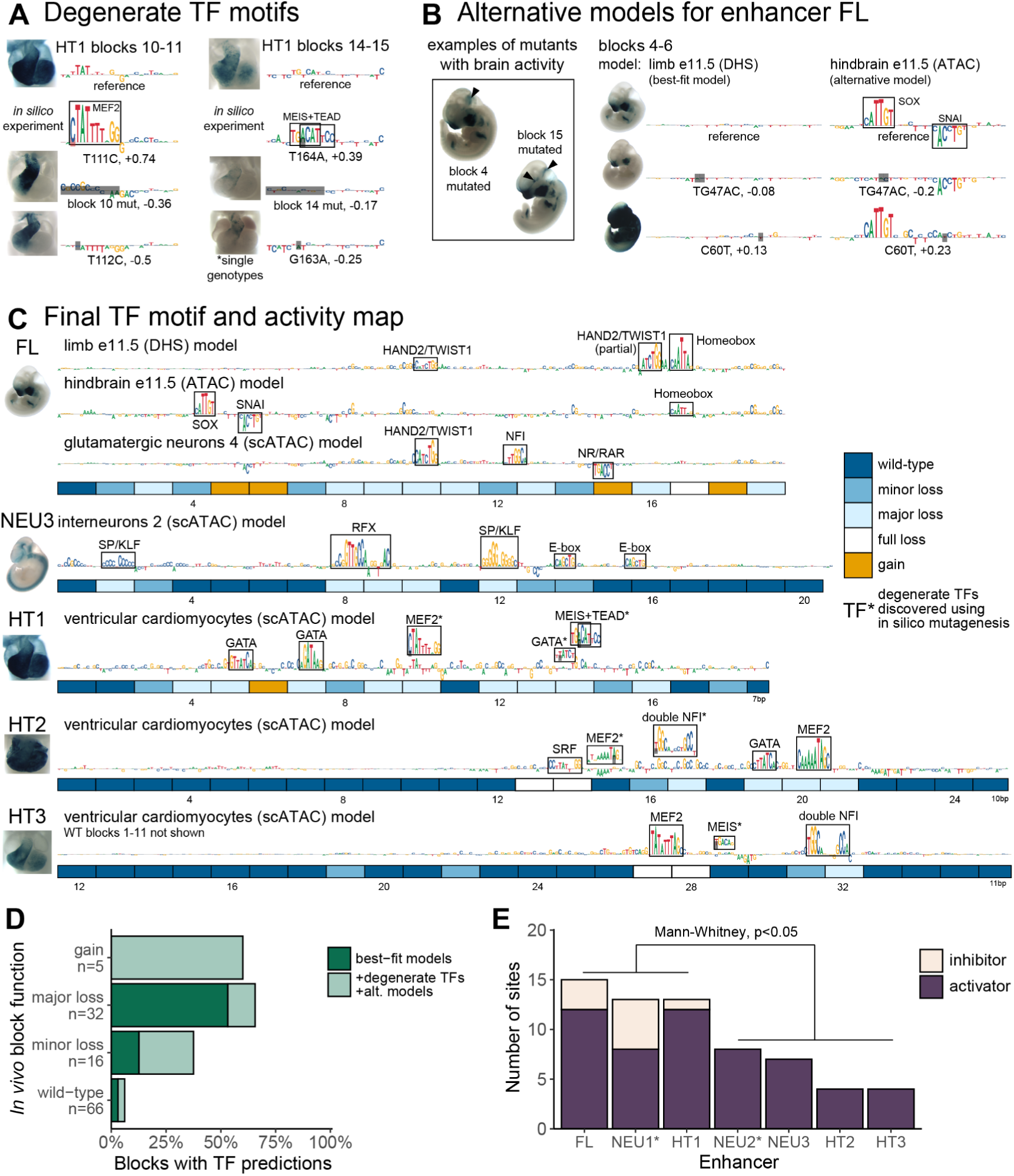
Refined map of binding motifs and enhancer activity. (A) Discovery of additional sites through *in silico* mutagenesis and validation. Also see Supplementary Figure 5A and B. (B) Examples of block mutants with gain of brain activity and additional motifs discovered using alternative FL models trained on neuronal datasets. Black arrowheads indicate gain of function. Also see Supplementary Figure 5C. (C) Final TF binding motif and activity map. Includes motifs discovered using alternative models (element FL) and degenerate motifs (marked with asterisks; elements HT1, HT2 and HT3). (D) Fraction of blocks with motif predictions, by experimentally determined function. Major loss includes full loss. (E) Number of activator and inhibitor sites as estimated from experimental data alone (marked with asterisk; NEU1 and NEU2) or from experimental data combined with model motif predictions (FL, NEU3, HT1-3), by enhancer (Methods, Supplementary Figure 5D for visual guide).

We also explored if the sensitivity of detecting functional sites can be further increased by combining models derived from multiple training sets representing different relevant tissues. We tested this paradigm using face and limb enhancer FL, which showed a striking increase in activity in the brain in several block mutation experiments, suggesting latent neuronal activities that could potentially be studied using models derived from brain tissues containing many different types of neuronal cells (Figure 3B, left). Indeed, using an alternative model derived from e11.5 hindbrain ATAC-seq data, we observed two strong binding site motif predictions for activator SOX and repressor SNAI that were not apparent in the best-fit limb model (Figure 3B, right). A targeted 2-basepair mutation of the SOX motif resulted in loss of *in vivo* function, whereas a targeted single-basepair mutation in the repressive SNAI motif caused a major gain of *in vivo* function (Figure 3B). Using an additional model derived from glutamatergic neurons, we observed two more sites, including a repressive NR/RAR motif located in a sequence block that causes a gain of activity when mutated (Supplementary Figure 5C). Together, the use of two alternative models identified four additional binding motifs in enhancer FL, thereby providing mechanistic explanations for the observed *in vivo* activity changes.

The combined use of *in silico* saturation mutagenesis and alternative models (Figure 3C) predicted TF motifs in 30 of the 53 blocks that showed altered *in vivo* activity upon mutation, increasing sensitivity to 59% compared to 36% based on best-fit models alone. Despite this substantially improved sensitivity, we observed only a minor reduction in positive predictive value, from 14/15 (93%) to 22/25 (88%) of predicted functional sites showing altered *in vivo* activity. Blocks classified to cause a major loss of function when mutated had a predicted TF site more often than those causing minor loss of function (66% vs 38%), although the difference was not statistically significant (p=0.06, Fisher’s Exact Test; Figure 3D).

Combining the results of block mutagenesis and open chromatin model predictions also offers an opportunity to examine the overall complexity of individual enhancers by estimating the total number of functional sites (Methods, Supplementary Figure 5D). We observed that the seven enhancers examined had between 4 and 15 functionally relevant sites (average: 9; Figure 3F). Enhancers that contained blocks which, when mutagenized, caused a gain of function, had the highest number of sites (13-17 sites total; FL, NEU1, HT1; p<0.05, Mann-Whitney U-test). Taken together, these results show how combining large-scale *in vivo* mutagenesis, epigenomic data, and predictive modeling can elucidate the functional landscape of *in vivo* enhancers at base-pair resolution.

### Response to Mutations Reveals Regulatory Modes

The complex spatial activity patterns of enhancers, which frequently include multiple developmental tissues and cell types, represent an additional hurdle for relating enhancer sequence content to *in vivo* function. We explored whether the results of our mutagenesis screen can be used to disentangle the relationship between sequence features within an enhancer and tissue-specific activities.

First, we examined the impact of different mutations on the *in vivo* activity of enhancer HT1 in different subregions of the developing heart. The reference allele of HT1 showed strong activity in the outflow tract and both ventricles, along with weaker activity in the atria (Figure 4A). We scored the activity changes observed in each of these four cardiac subregions for each single-block mutagenesis allele in comparison to the reference allele (Figure 4A). We observed that activity in the atria and left ventricle was typically more severely affected than activity in the right ventricle and the outflow tract. Extending this analysis to include constructs with multiple mutated blocks, and sorting them by the overall severity of the observed changes (Figure 4B) revealed a graded response in which atrial expression was most susceptible to mutations, followed by left ventricle, right ventricle, and outflow tract. We did not observe any cases in which outflow tract or right ventricle expression was affected in the absence of changes to left ventricle or atrial activity. This suggests that functional sites within enhancer HT1 predominantly do not drive expression in specific subregions of the heart, but contribute to an overall pattern in a graded fashion. A similarly graded response was observed for enhancer NEU2 (Supplementary Figure 7B).

**Figure 4.**
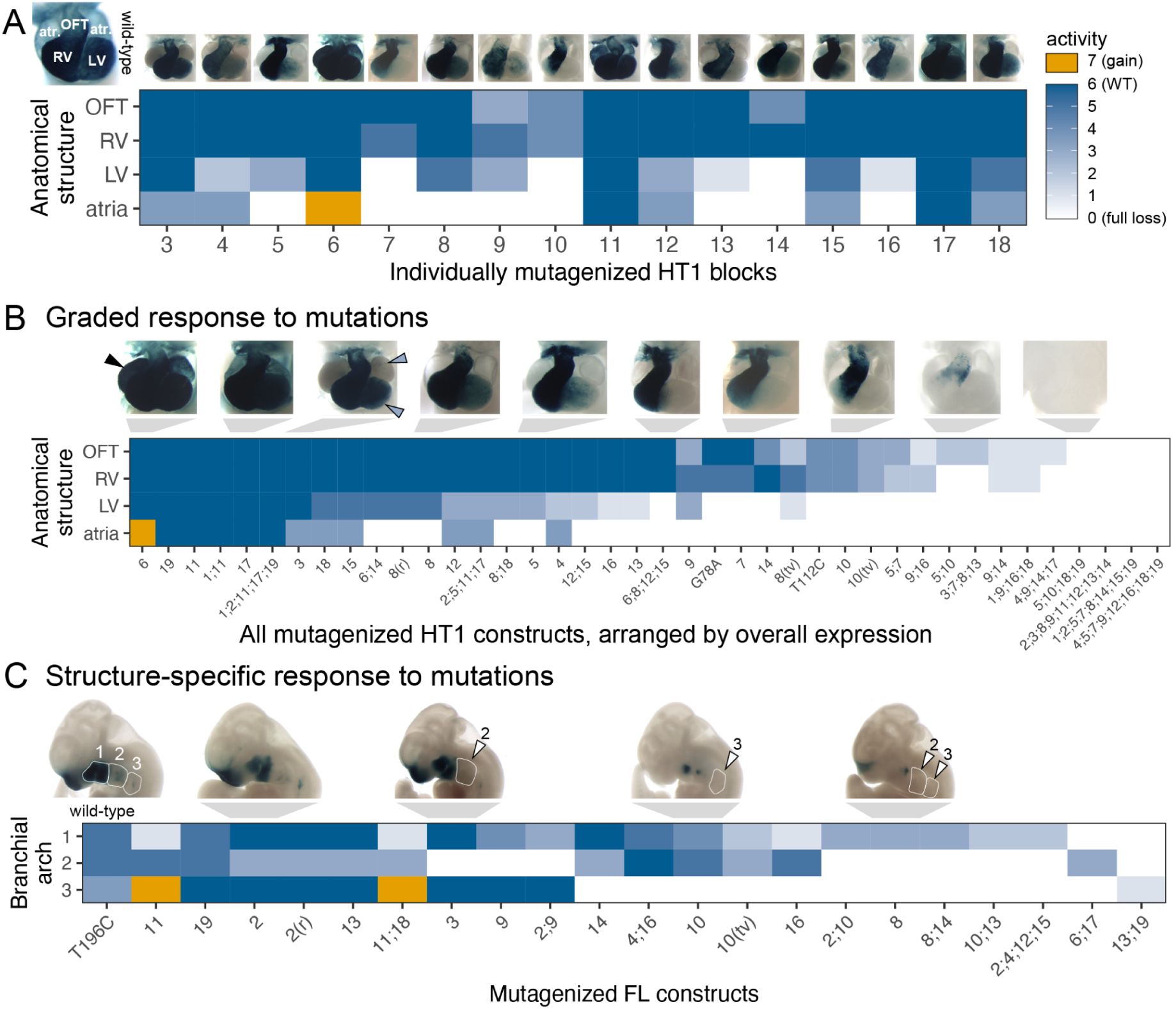
Patterns of multi-tissue *in vivo* responses to mutations. (A) Activity of single block mutants of enhancer HT1, scored across four cardiac substructures. Flanking wild-type blocks not shown. (B) Activity all mutated HT1 constructs, scored across four cardiac substructures, arranged by overall expression (Methods). (C) Activity of mutated FL constructs, scored across three branchial arches. Arranged by structure-specific full loss of function. Only mutants with partial loss of function in one of the arches were included. OFT = outflow tract, LV = left ventricle, RV = right ventricle, atr. = atrium, (r) = random scrambling mutagenesis, (tv) = GC content preserving transversion mutagenesis, 1;11 = combinatorial mutagenesis of blocks 1 and 11, A190G = 1bp A to T mutation at position 190. Arrowheads: black = gain of function, blue = minor loss, white = full loss. Also see Supplementary Figure 7.

Next, we examined enhancer FL, which shows more complex expression changes, performing the same structure-specific annotations (Supplementary Figure 7A). Focusing on expression in the first, second, and third branchial arch, we observed structure-specific activity changes associated with distinct subsets of mutations (Figure 4C). For example, mutations of blocks 3 or 9 selectively abolished expression in branchial arch 2 while maintaining activity in branchial arches 1 and 3. In contrast, mutations of blocks 10 or 16 selectively abolished expression in branchial arch 3. These results show that distinct aspects of the complex *in vivo* activity pattern of enhancer FL require different functional subregions of the enhancer. A similar structure-specific response to mutations was observed for enhancer NEU1 (Supplementary Figure 7C).

In contrast to these structure-specific effects of mutations affecting branchial arch activity, some other tissues in which enhancer FL is active exhibited graded responses more similar to HT1 and NEU1. In particular, all mutations that caused a full loss of activity in any facial substructure also caused loss of limb activity (consistent with shared developmental signaling in these tissues^26^). Conversely, nearly all incomplete loss mutants (27/29) retained some activity in branchial arch 1 (Supplementary Figure 7A). These findings indicate that all functional sites within enhancer FL contribute to limb and branchial arch 1 activity in a graded fashion, while some functional sites are specifically required for activity in either branchial arch 2 or branchial arch 3.

In conclusion, the results of our large-scale *in vivo* enhancer mutagenesis highlight two distinct modes by which mutations can affect the activity of enhancers with complex, multi-tissue activity patterns. The more commonly observed mode is a graded response of structures to mutations, with some structures being overall more sensitive to mutations than others. In a second strictly structure-specific mode, distinct mutations affect activity in distinct substructures independently. As illustrated by enhancer FL, these modes are not a general property of a given enhancer, but can co-occur within the same enhancer, applying to different aspects of the complex activity pattern.

### Paired Block Mutagenesis Demonstrates Pervasive Additive Logic

The severity of activity changes in enhancers generally increased with the number of introduced mutations (see, e.g., Figure 4B). However, this observation does not immediately reveal the functional impacts expected from compound mutations that affect more than one functional sequence block. Under a simple model of enhancer function, individual sites within the enhancer contribute to the enhancer’s overall regulatory activity in an additive fashion. Consequently, it is expected that combinations of mutations cause additive *in vivo* activity changes that reflect those observed in the constituent single-block mutagenesis experiments. However, more complex modes of functional intra-enhancer interaction resulting from compensatory or synergistic functional interactions between sites are also conceivable^27–30^. To examine the prevalence of such complex interactions in human *in vivo* enhancers, we systematically compared how mutagenesis of single 12bp blocks or pairs of such blocks affected *in vivo* activity. We only studied pairs separated by at least one block to avoid potentially confounding gain-of-binding events at the boundary of adjacent blocks and to exclude short-distance, homo- and heterodimer TF interactions. Under an additive model of function, we expected combining two loss-of-function mutations to result in a more pronounced loss. Any other outcome would indicate deviation from the additive model (Figure 5A; Supplementary Figure 8A).

**Figure 5.**
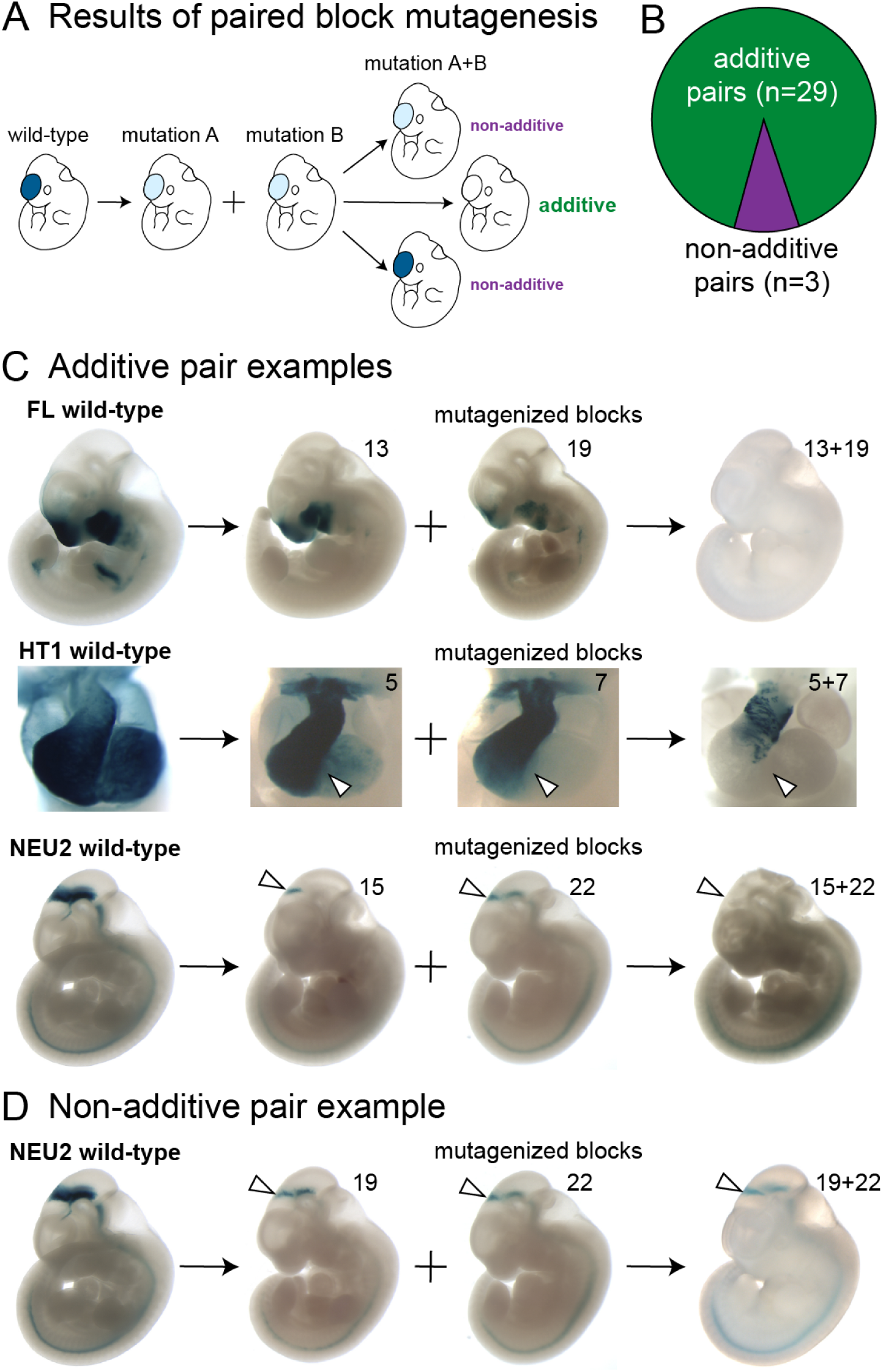
Comparison of individual and paired block mutations. (A) Classification of outcomes of paired block mutagenesis. A combination of two loss-of-function mutations resulting in a more pronounced loss is considered additive, while any other outcome is classified as non-additive (also see Supplementary Figure 8A). (B) Distribution of additive and non-additive outcomes of paired block mutagenesis. (C) Examples of additive pairs. (D) An example of non-additive pair. White arrowheads highlight structures of interest (see main text). Also see Supplementary Figure 8.

We examined 32 pairs of blocks and found 29 (90%) to have patterns consistent with the additive model (Figure 5B). For example, combined mutation of blocks 13 and 19 of enhancer FL resulted in full loss of function, while mutagenesis of either of these blocks in isolation led to only incomplete reduction in enhancer activity (Figure 5C). A similar additive effect was observed for HT1 blocks 5 and 7, as well as NEU1 blocks 15 and 22, for which paired block mutation caused more severe loss than either of the individual block mutations (Figure 5C; see Supplementary Figure 8B for additional examples). As a contrasting example of non-additive changes in function of a compound mutagenesis construct, partial loss of midbrain activity caused by mutagenizing NEU1 blocks 19 and 22 together was highly similar to the effect of mutagenizing either of the blocks in isolation (Figure 5D; see Supplementary Figure 8C for remaining non-additive pairs). Taken together, our results demonstrate that most functional sites within human *in vivo* enhancers contribute to overall regulatory activity of the enhancer in an additive manner. More complex functional interactions between sites within an enhancer can occur, but are rare.

## Discussion

Over the past decade, dramatically improved maps of the transcriptional enhancers orchestrating human genome function have emerged from genome-wide mapping efforts in hundreds of mammalian tissues and cell types^31–34^. In sharp contrast, our understanding of the genomic code for how individual enhancers direct gene expression *in vivo* remains cursory. This knowledge gap currently prevents accurate predictions of how a given mutation within an enhancer will impact its *in vivo* function. To develop a systematic and robust data foundation for gaining insight into this relationship, we performed comprehensive *in vivo* mutagenesis mapping of multiple human developmental enhancers with different tissue specificities, leveraging mouse genome editing to generate and analyze more than 1,700 independent transgenic mouse embryos. Our studies revealed a diversity of functional site arrangements within these enhancers, enabled the identification of machine learning models for prediction of functional binding motifs at basepair resolution, identified strategies to improve the sensitivity of machine learning models, described complementary modes of multi-tissue activity, and established an additive model as the predominant mode of functional site interactions.

Systematic block mutagenesis of seven *in vivo* enhancers showed that all had a complex functional architecture, with sites required for normal activity spread across hundreds of basepairs, and revealing pronounced differences in overall sensitivity to mutations (Figure 1). Three enhancers could be completely inactivated by mutagenesis of a single “Achilles’ heel” block. Conversely, three enhancers contained blocks which, when mutagenized, led to gains of function. In an example of extremely high density of functional sites, no single core block of enhancer FL could be mutagenized without affecting its *in vivo* activity. In contrast, in an example of low density of functional sites, the majority of blocks in the functional core of enhancer HT3 could be mutated without impact on the observed *in vivo* activity. Given the spectrum of density of functional sites observed across the enhancers studied here, we speculate that even more robust enhancers, in which no individual block mutation leads to major loss of function, may exist.

Predictive modeling greatly complemented our experimental survey, allowing us to interpret the results of block mutagenesis at basepair resolution, with considerable sensitivity and high positive predictive value (Figure 2 and Figure 3). Systematic comparison of models against experimental data from *in vivo* block mutagenesis enabled the selection of best-fit models for individual enhancers. We found that the models trained directly on tissue-specific open chromatin signal predicted coherent, tissue-appropriate sets of binding motifs. The resulting high-confidence predictions enabled the targeted experimental verification of functionally relevant nucleotides within each block, highlighting a powerful computationally guided strategy for the interpretation of human pathogenic mutations and evolutionary divergence at enhancers across species.

By applying machine learning models to *in silico-*mutated enhancer sequences, we uncovered additional, degenerate TF motifs that could not be detected in reference sequences, thereby further increasing model sensitivity (Figure 3A, Supplementary Figure 5A). Notably, despite their low contribution scores in the context of the wildtype enhancer, we showed experimentally that these motifs contribute to the *in vivo* function of the respective enhancers. This observation aligns with the notion that suboptimal, lower-affinity TF binding sites in enhancers contribute to tissue-specific activities^35–38^. Application of machine learning models to *in silico-*mutated enhancer sequences offers an effective and scalable approach for the systematic discovery of such sites in other enhancers.

Three out of seven enhancers in our study harbored blocks that, when mutagenized, caused gains of activity, either in tissues in which the wildtype allele is inactive or quantitatively increasing activity in a tissue in which the wildtype allele is active. Such gains of function suggest the presence of repressive binding sites within these blocks, resulting in tissue-specific derepression upon mutagenesis. Generally, enhancers that included gain-of-function blocks also appeared to have overall more functional sites than enhancers that contained only blocks that caused loss of function when mutated (Figure 3E). The two enhancers containing multiple gain-of-function blocks (FL and NEU1) also had the clearest examples of mutations acting in a structure-specific manner (Figure 4A). This suggests that the activity in different tissues is enabled by the interplay of activating and repressive sites, which is consistent with observations of activator-repressor logic in other developmental enhancers^29,39,40^.

The complexity of functional impacts of mutations across tissues stresses the importance of studying human enhancers using whole-organism, multi-tissue experimental paradigms. For example, several of the gain-of-function activity changes we observed in face/limb enhancer FL appeared in unrelated organ systems, such as the heart and nervous tissues (Supplementary Figure 2B, Figure 3B). This aligns with our observation that some of the functional motifs for enhancer FL were not detected by machine learning models trained only on tissues in which the reference enhancer was predominantly active, namely face and limb (Figure 3C, Supplementary Figure 5C). It would be challenging to capture such mutation-induced ectopic activity in unexpected tissues even in complex *in vitro* systems. Our findings imply that interpretation of human non-coding variation and regulatory evolution, as well as designing safe, tissue-specific gene therapies will require a multi-tissue, *in vivo* approach, taking into account a possibility of ectopic activation from as little as a single basepair mutation (Figure 3B).

Systematic mutagenesis also provided insight into the relationship between individual sequence features of enhancers and their respective function in directing complex activity patterns that include multiple tissues or anatomical regions (Figure 4). In particular, we observed that most mutations caused a quantitative reduction in activity relative to the wild-type baseline activity across all tissues. Since baseline activity may vary across tissues, this resulted in a general graded reduction in activity across tissues. However, we also observed several cases in which mutations affected *in vivo* activity selectively in individual anatomical structures, implying that the corresponding wildtype sequence feature interacts with TFs with spatially restricted expression.

Combining mutations in pairs of blocks allowed us to examine the possible presence of functional interactions between sites. We observed examples of additive effects on enhancer function, where the combined mutations resulted in additive *in vivo* activity changes, as well as non-additive effects. In 90% of cases, we found a simple additive pattern, suggesting that additive logic is the predominant mode in human developmental enhancers (Figure 5). The non-additive cases we identified may represent opposing or interfering effects of two TFs. Alternatively, they may be a special case of additive logic, in which block mutations simultaneously lead to a loss of activity in one cell type and a gain of activity in another cell type in the same anatomical structure. The effect of combining such block mutations may appear to be non-additive. Identifying the underlying TFs will help design experiments to interpret these observations.

In conclusion, our comprehensive mutagenesis survey of human *in vivo* enhancers revealed many facets of within-enhancer regulatory logic, in particular pertaining to activator-repressor paradigm, multi-tissue expression and applicability of predictive modeling. These findings provide a foundation for the interpretation of human non-coding variation, changes of enhancer activity across evolution, and will aid in the design of synthetic enhancers for biotechnological and therapeutic purposes.

## Supplementary Figures

**Supplementary Figure 1.**
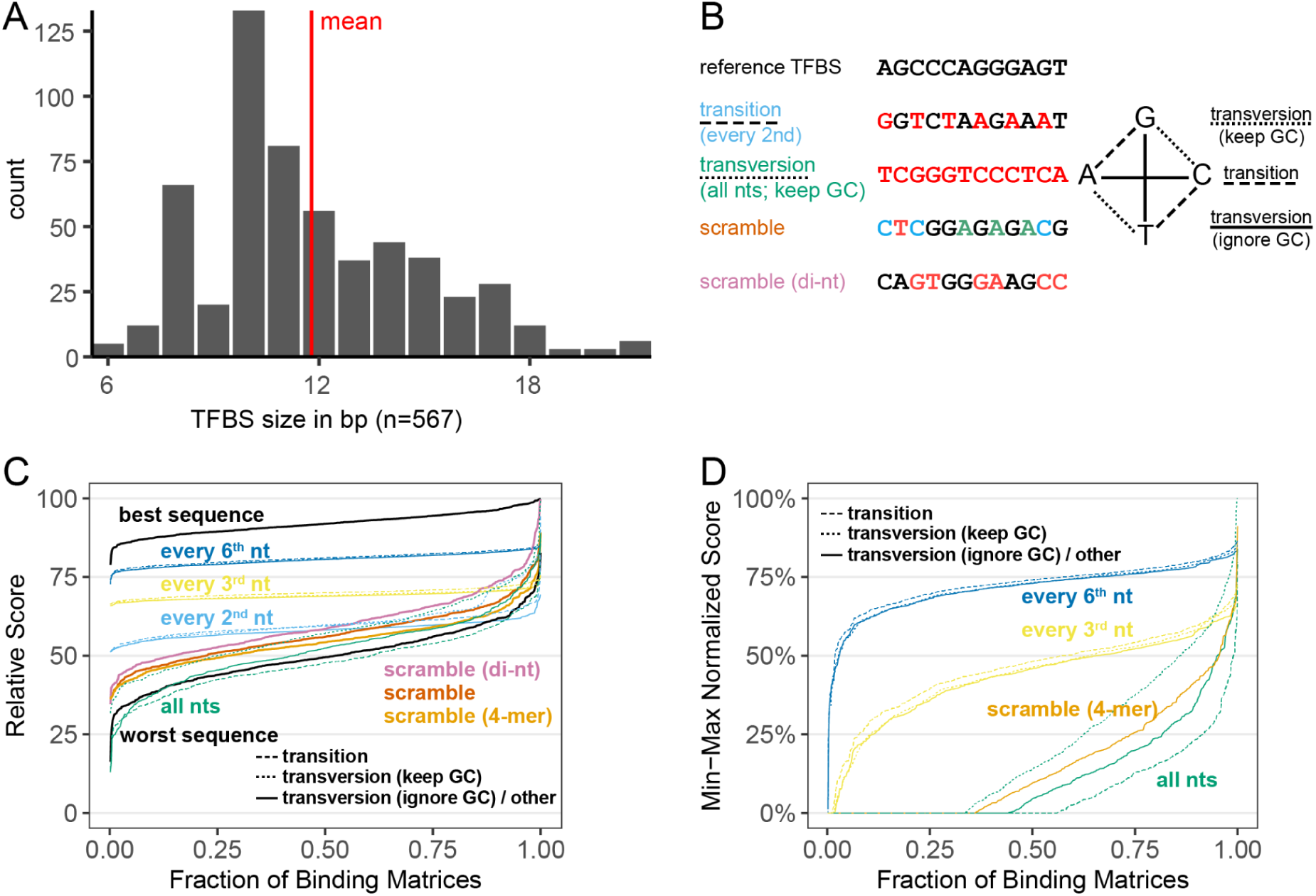
Choice of mutagenesis strategy. (A) Size distribution of all JASPAR TF binding motifs. (B) Visualization of *in silico* mutagenesis schemes. (C) Relative score of matches between original TF PWM and mutagenized sequence. (D) Match score min-max normalized to that of best and worst sequence for a given TF PWM. See text for details. Observations are ordered on x-axis by score, so each position does not correspond to the same TF PWM.

**Supplementary Figure 2.**
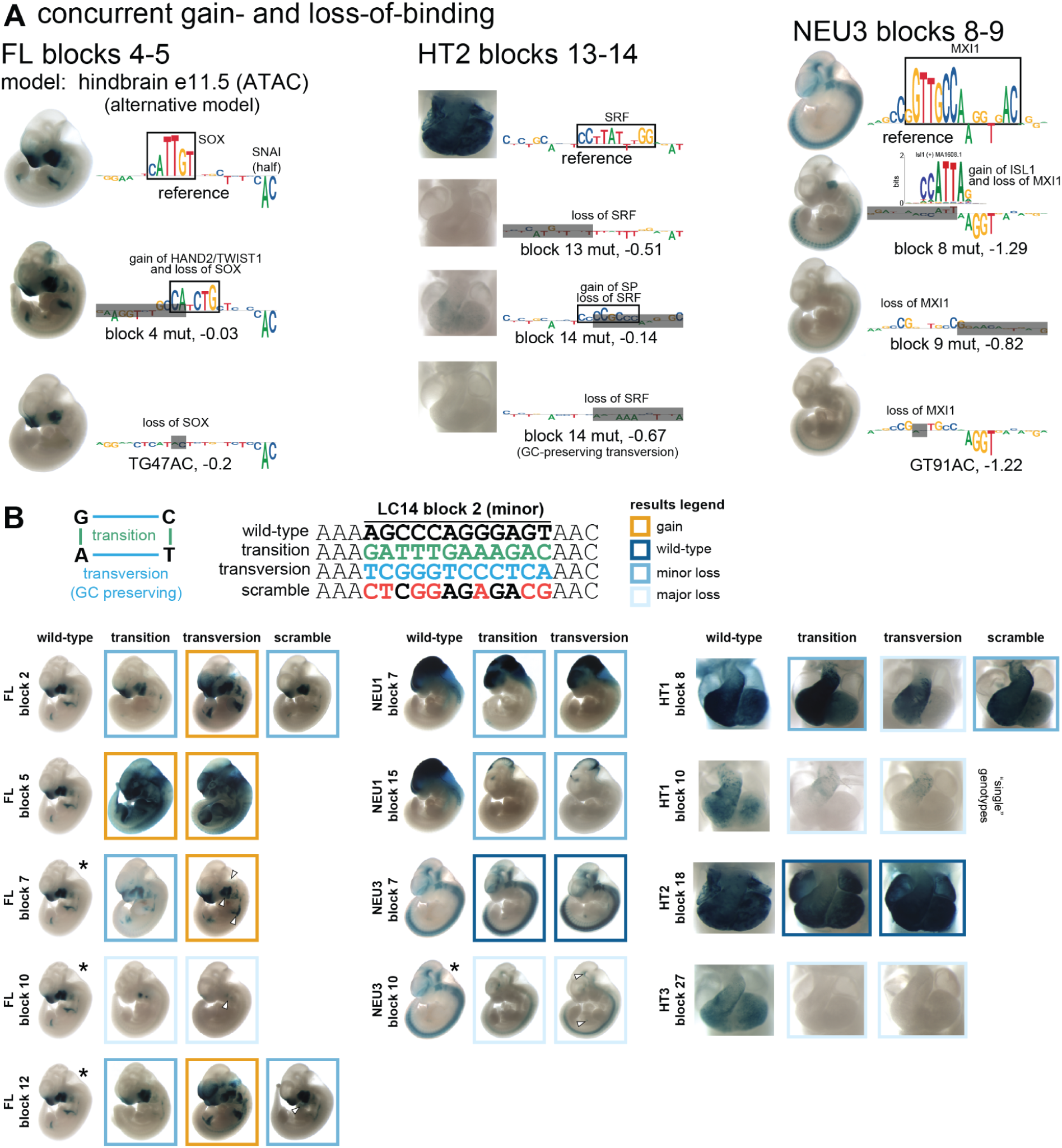
Validation of transition mutagenesis scheme. (A) Three blocks with suspected gain-of-binding events or mismatch between adjacent blocks overlapping the same predicted binding motif were tested using alternative mutagenesis scheme or targeted 2bp mutations. In all cases, a result confirming gain-of-binding was obtained. (B) Unbiased testing using alternative mutagenesis schemes. Blocks were mutagenized using both a deterministic transition scheme (default for this study) and a GC-preserving transversion scheme, with selected blocks also mutagenized through random scrambling. Tandem embryos are displayed, except when indicated otherwise (see Methods for genotype definitions). White arrowheads indicate regions in which results of alternative mutagenesis mismatch those of transition mutagenesis (blocks marked with asterisk). See Supplementary Note 2 for details. Related to Figure 1.

**Supplementary Figure 3.**
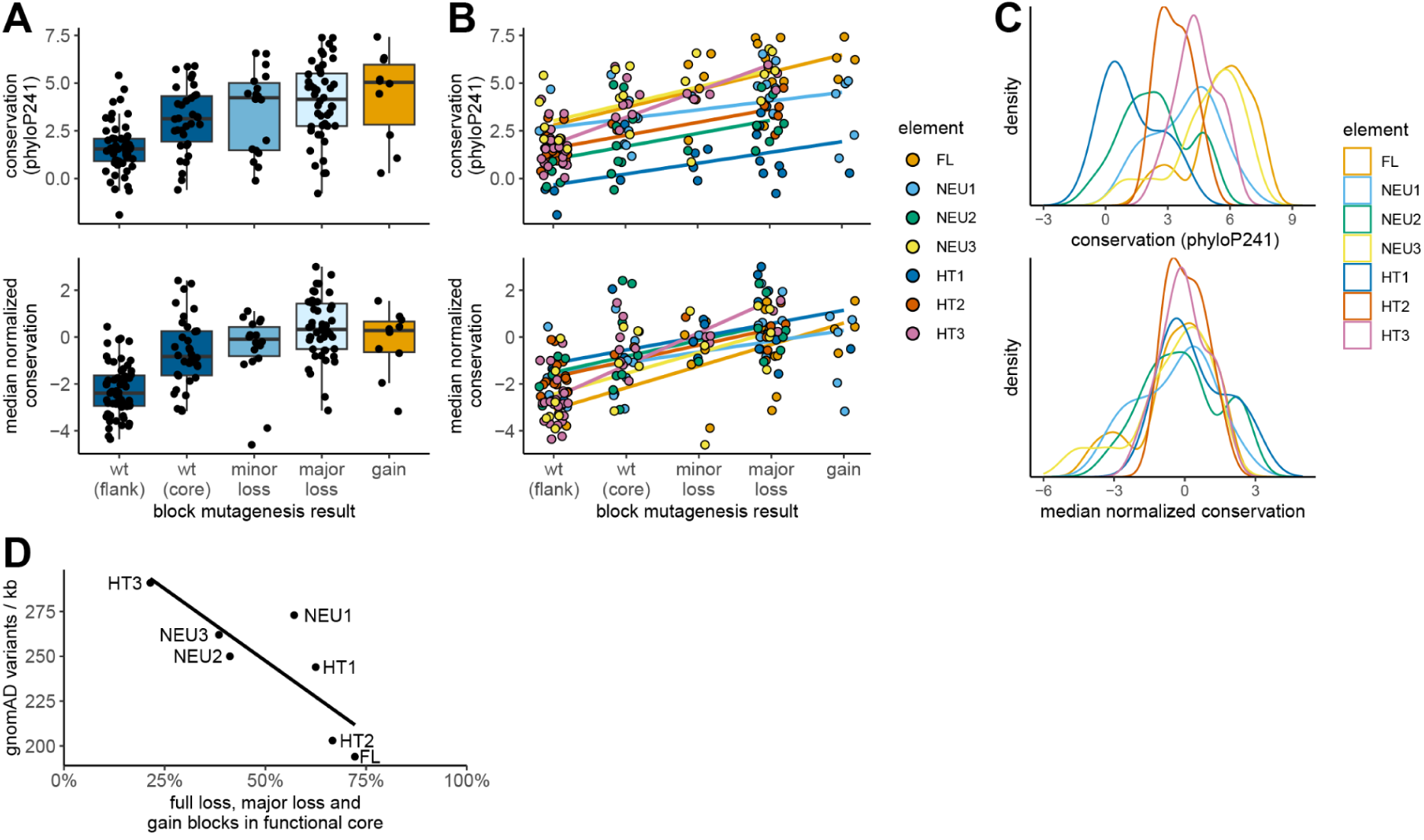
Conservation score normalization and analysis including flanking wild-type blocks. (A) Conservation score boxplots by block mutagenesis result. (B) Same as A, but colored by enhancer. Linear regression line is added. (C) Density of conservation scores, colored by enhancer. Each dot in A and B is a 12bp block (N=167). Top panels use raw mammalian conservation score (phyloP241), bottom panels use raw score normalized for median of functional core (per enhancer). Minor loss, major loss and gain blocks were each more conserved, after median normalization, than either wild-type flanking blocks or all wild-type blocks combined. Only major loss blocks were more conserved than wild-type core blocks (FDR<0.05, 9 comparisons, 7 significant). (D) Correlation between density of gnomAD variants and fraction of functional blocks in functional core (Pearson R^2^=68%, p-value<0.05). Related to Figure 1.

**Supplementary Figure 4.**
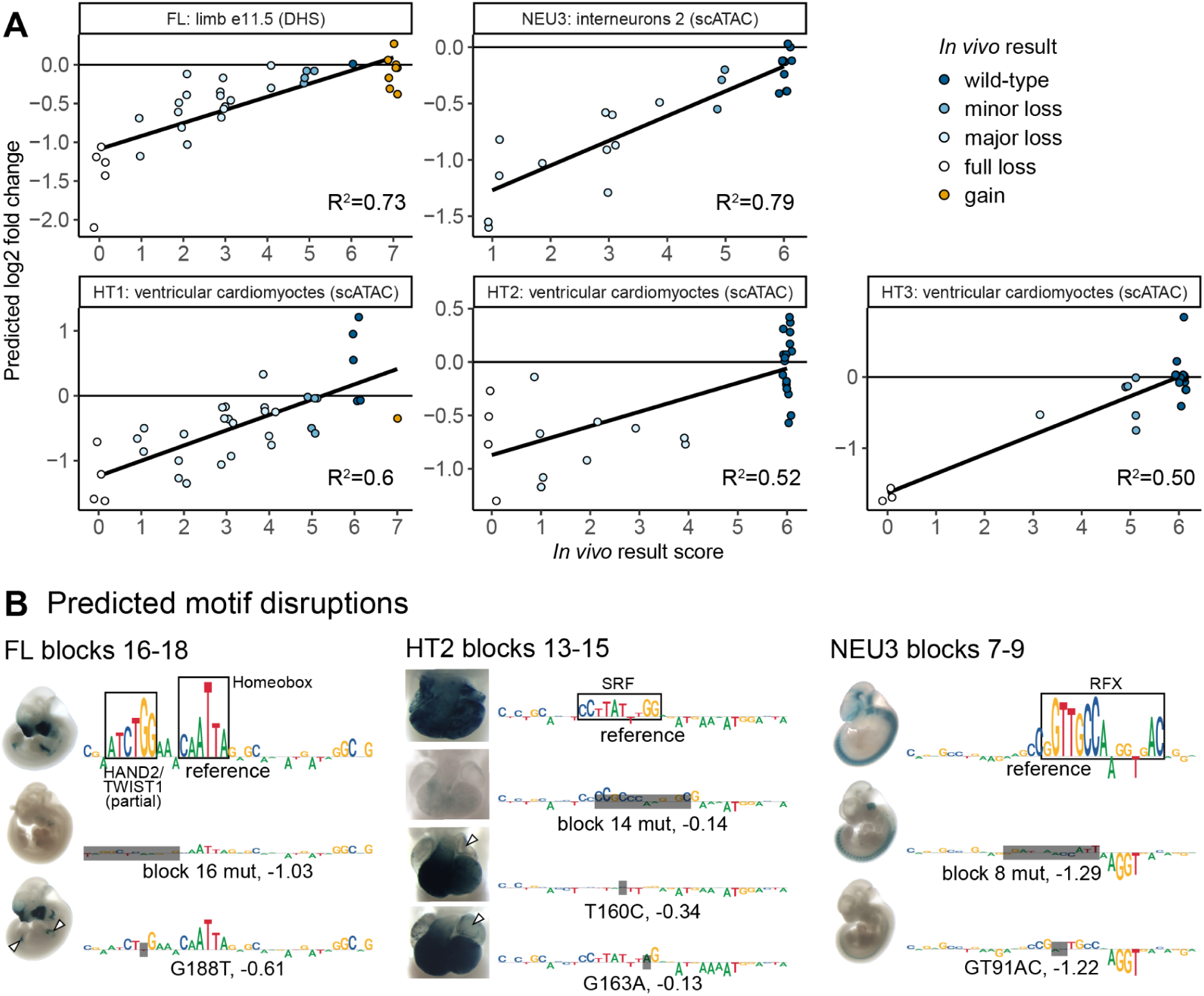
Machine learning model selection and validation. (A) Correlations between model predictions and *in vivo* results. Dots = mutagenized constructs. Black fit line is linear regression. R^2^ is Spearman correlation. (B) Remaining predicted motif disruptions. Related to Figure 2.

**Supplementary Figure 5.**
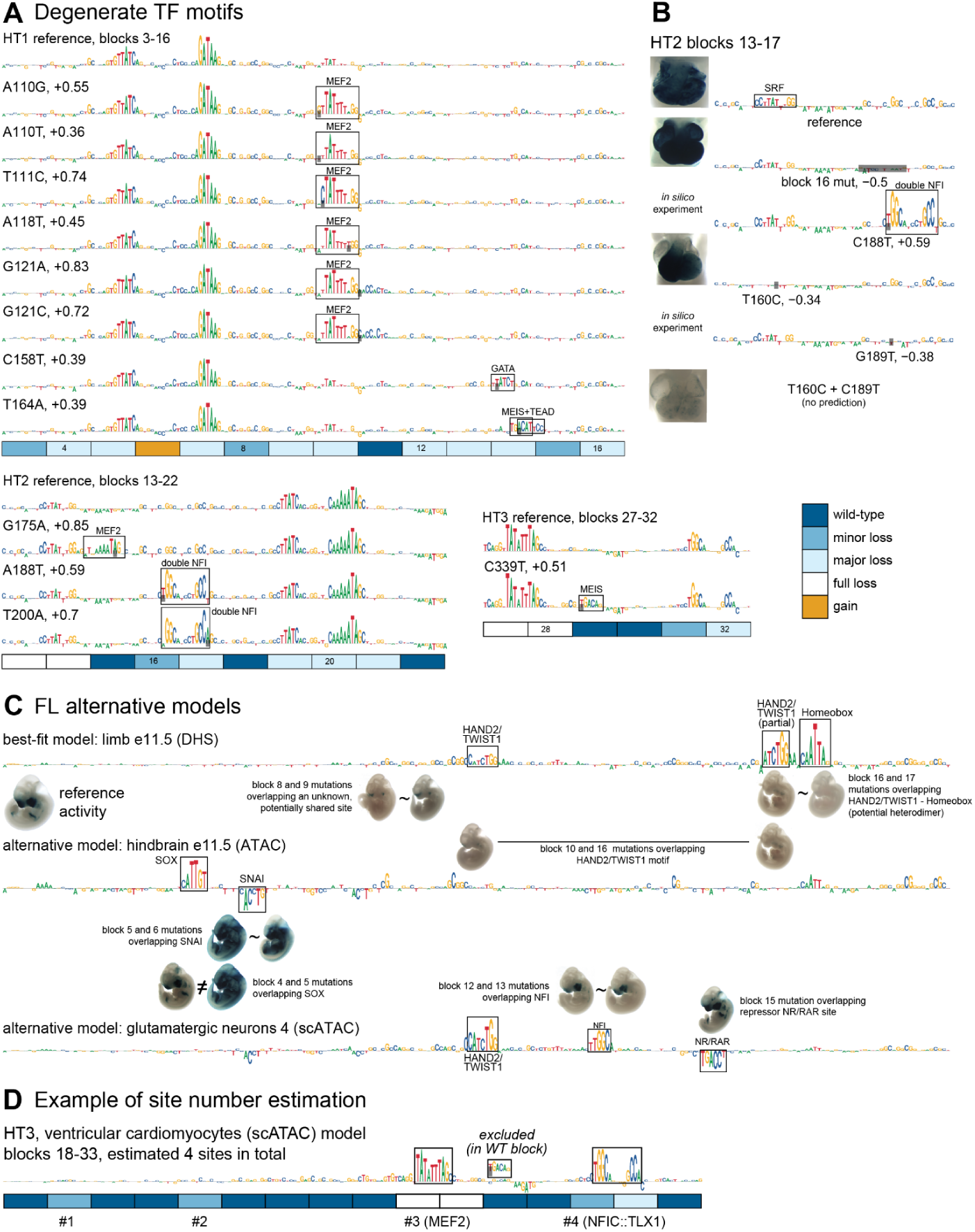
Degenerate TFs and alternative models. (A) Contribution score tracks for wild-type sequences and *in silico* mutated constructs which were predicted both to increase the open chromatin signal by at least 25% (log2 fold change > 0.32) and to feature a novel cluster of high, positive scores. Three of six discovered sites were validated experimentally. Two of the unverified sites that overlapped wild-type blocks were classified as false positive predictions (MEF2 in HT2 and MEIS in HT3). (B) Validation of double NFI site predicted in blocks 16-17 of enhancer HT2 by *in silico* saturation mutagenesis. Combined 1bp mutations in SRF site (T160C) and in the predicted double NFI site (C189T) led to a more pronounced loss of function than SRF mutation alone. This validated the double NFI site and led to reassessment of block 16 as (at least) minor loss. Supplement to Figure 3A. (C) Exploration of alternative models for enhancer FL. Block mutations overlapping the same binding motif show very similar activity impacts, with exception of block 4 and 5 (see Supplementary Figure 2 and Supplementary Note 2). (D) Example of total site count for enhancer HT3 (all functional blocks shown). Total site count = all predicted sites – predicted sites in wild-type blocks + blocks without site predictions (4=3-1+2 in this case).

**Supplementary Figure 6.**
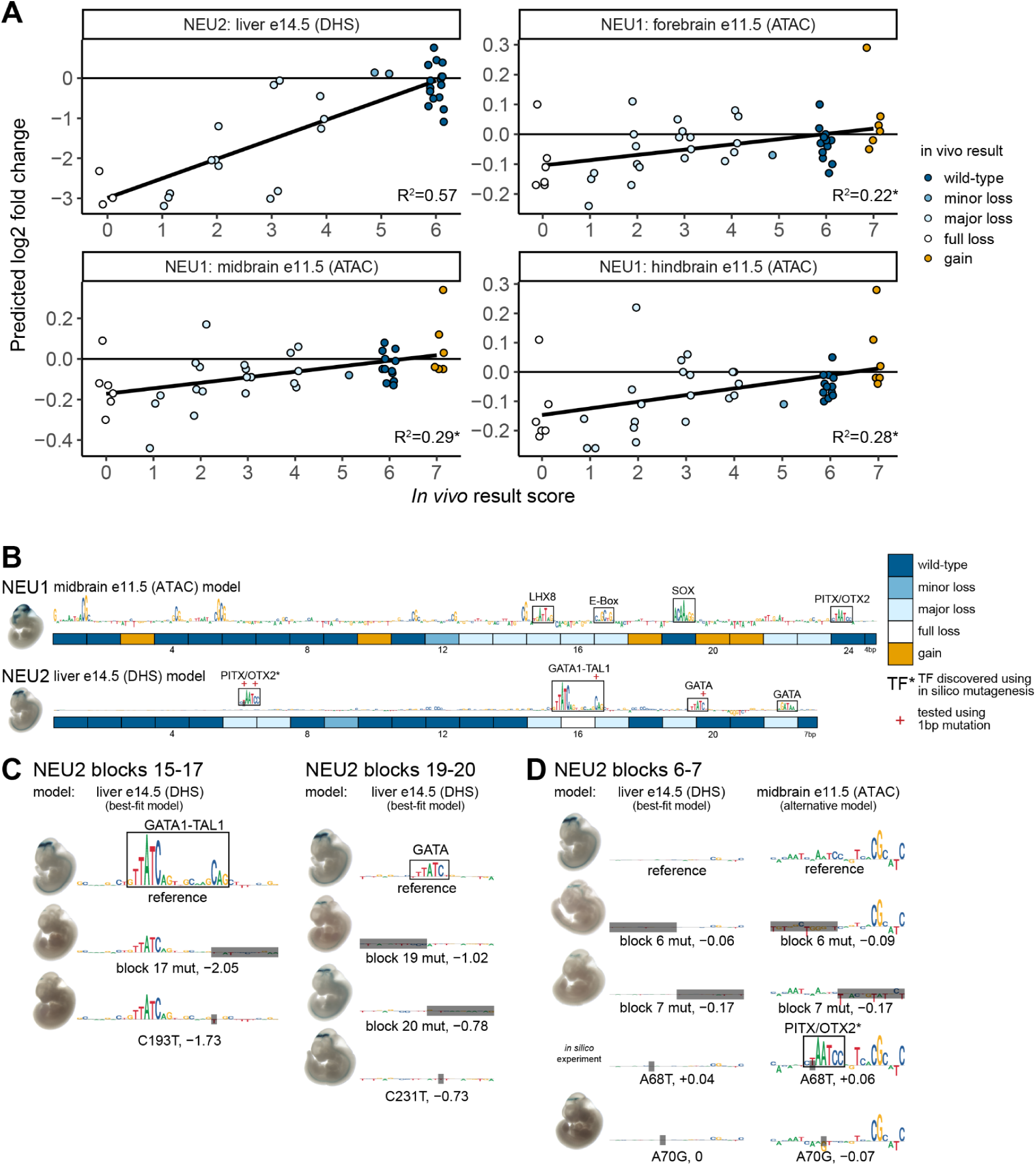
Rejected best-fit machine learning models for enhancers NEU1 and NEU2. (A) Correlations between model predictions and *in vivo* results. Dots = mutagenized constructs. Black fit line is linear regression. R^2^ is Spearman correlation. Asterisk = non-significant (FDR>0.01). (B) Final TF binding motif and activity map including verified binding motifs discovered through *in silico* saturation mutagenesis (NEU2 PITX/OTX2 site marked with asterisks). (C) Predicted motif disruptions. Note that validation of the GATA motif in blocks 19-20 did not succeed. (D) Discovery and validation of an additional PITX/OTX2 site in enhancer NEU2.

**Supplementary Figure 7.**
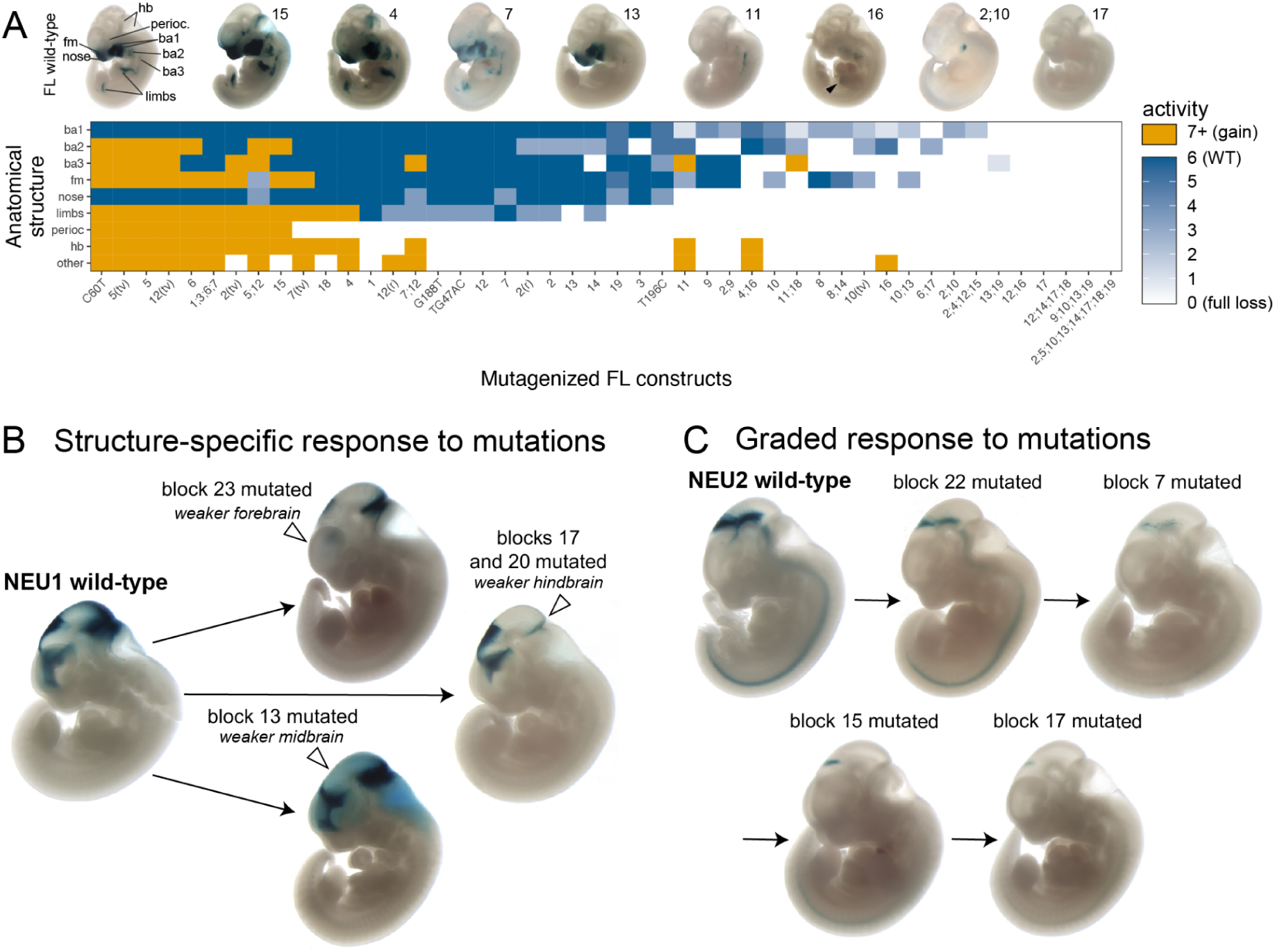
Additional examples of multi-tissue responses to mutations. (A) Illustration of paired block mutagenesis outcomes for all possible combinations of loss and gain mutations. Bars represent ranges of possible outcomes that would be classified as additive or non-additive. Redundant is a special case of non-additive in which combined mutagenesis of two blocks resulted in an outcome exactly as severe as the most severe of individual block outcomes. (B) Additional additive pair examples. (C) Remaining three non-additive pairs. White arrowheads indicate loss of function. Black arrowhead indicates gain of function. Related to Figure 4.

**Supplementary Figure 8.**
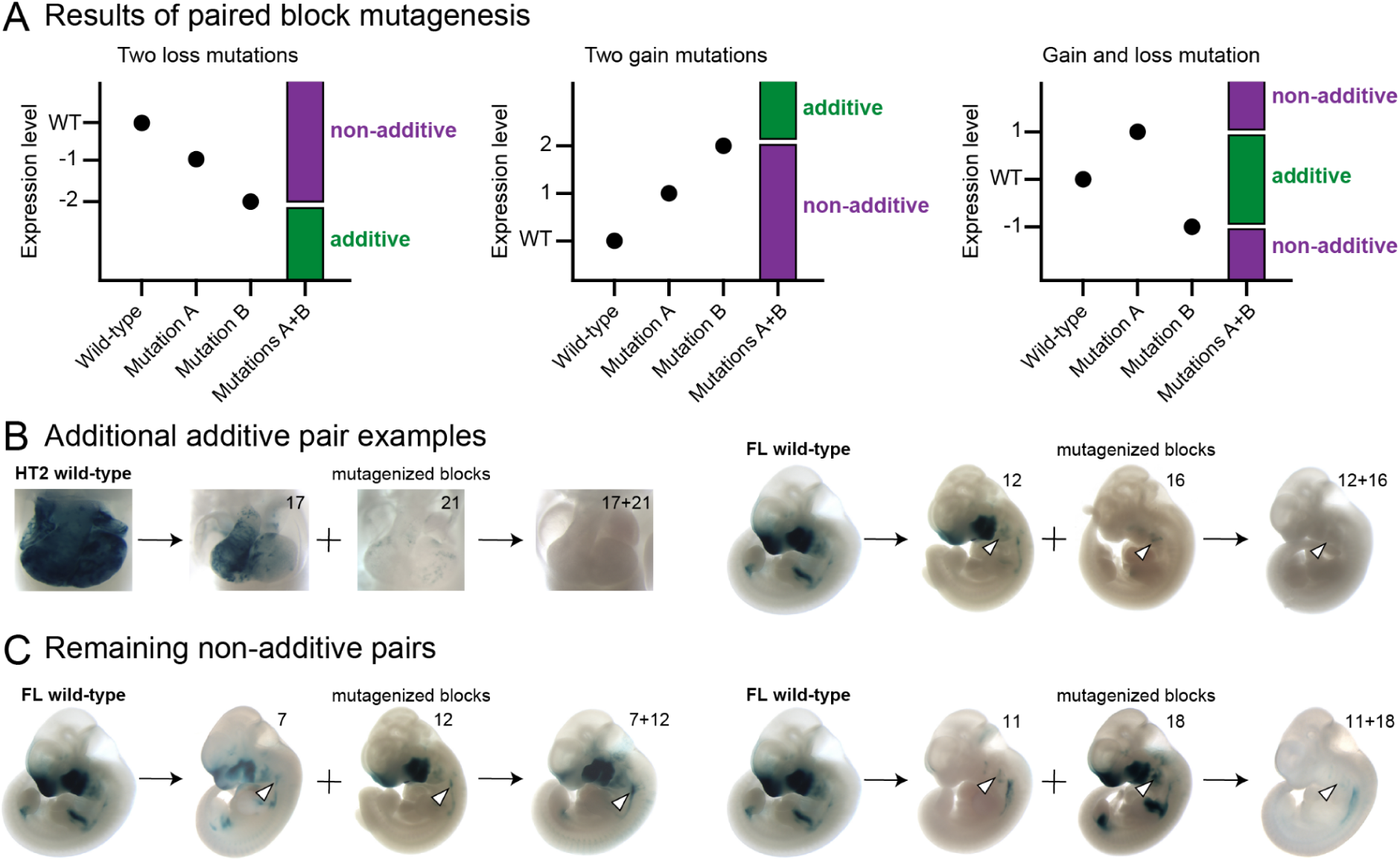
Additional results of paired block mutagenesis. (A) Illustration of paired block mutagenesis outcomes for all possible combinations of loss and gain mutations. Bars represent ranges of possible outcomes that would be classified as additive or non-additive. (B) Additional additive pair examples. (C) Remaining non-additive pairs. Combined mutagenesis of enhancer FL blocks 7 and 12 resulted in higher branchial arch 3 activity, while no change in activity in these structures was observed in constructs with single block mutations (see also hindbrain activity). Combined mutagenesis of enhancer FL blocks 12 and 18 resulted in lower activity in branchial arch 2 compared to constructs mutated in block 12 only, while mutation of block 18 in isolation did not appreciably change the activity of this structure (compare also hindbrain activity). White arrowheads highlight structures of interest. Related to Figure 5.

## Supplementary Notes

### Supplementary Note 1: Deterministic transition mutations are the best strategy for eliminating existing TF binding motifs using block mutagenesis

In designing the mutagenesis scheme for this study, we aimed to achieve two goals - reduce the number of experiments necessary to comprehensively map the functional parts of chosen enhancers while retaining a reasonable sequence resolution and to avoid both false positives (calling a functional site in absence of function) and false negatives (calling a site that has function wild-type). We reasoned that mutagenizing sequences in blocks of 12bp, the average size of a TF PWM (Supplementary Figure 1A) strikes a good balance between the resolution and the throughput of the experiment. We speculated this would make it uncommon to deactivate two binding motifs by chance and if such contingency occurred, it would be rare enough to disambiguate using additional targeted mutations.

To choose a block mutagenesis scheme that prevents false negatives that may arise from TF binding motifs being accidentally recreated by mutations, we run an *in silico* experiment. We avoided indel schemes, reasoning that they could lead to changes in activity due to changes in spacing in between TF binding sites, which would make data interpretation difficult. We also avoided “homopolymer schemes”, e.g. replacing every basepair in a block with Ts, as that might substantially affect GC-content of the sequence and secondary DNA structure, leading to effects unrelated to changes in TF binding motifs. In the end, we chose to compare various deterministic and random scrambling strategies.

To validate our simulation, we mutagenized all, or every 6th, 3rd and 2nd nucleotide of TF binding site in JASPAR database^41^ and found that, as expected, the denser mutation schemes make it less likely for TF binding motifs sequence to retain match with the original PWM (Supplementary Figure 1B-D). We also tested three scramble schemes - simple randomization (’scramble’), randomization in blocks of two nucleotides often used in MPRA experiments (’scramble (di-nt)’) and a novel scramble scheme designed to randomize the sequence without recreating any of the 4-mers originally present (’scramble (4-mer)’). As expected, the di-nucleotide scramble was most likely to preserve TF binding motifs match, followed by random and 4-mer scramble. For non-scramble schemes, we tried all three possible deterministic mutations - transitions (A=G, C=T), GC-content preserving transversions (A=T, C=G) and transversions that did not preserve GC-content (A=C, T=G). In line with experimental results^42^, we found that the latter transversion scheme had a slight advantage over the other schemes when not mutagenizing every nucleotide. Surprisingly, when mutagenizing all nucleotides, transition scheme was much more potent than the two transversion schemes. Importantly, it was the most effective scheme across a range of TF binding motifs, more effective than 4-mer scramble. We select this scheme for our experiment.

Our simulations did not address the risk of false positives, ie mutagenesis creating a novel site resulting in a false functional call. We reasoned this will require both sacrificing the consistency of the deterministic scheme as well as an assumption that a good fraction of transcription factor binding motifs involved in the activity of all seven enhancers we have mutagenized are known. We decided that estimating this rate by employing alternative mutagenesis schemes post-factum to functional blocks detected by transition scheme is a better way of addressing this issue (Supplementary Figure 2).

### Supplementary Note 2: Transition scheme validation and gain-of-binding events

Transition block mutagenesis was primarily expected to lead to loss of existing binding motifs without creation of new binding motifs. To test this assumption, we used machine learning predictions from best-fit and alternative machine learning models to detect likely gain-of-binding events and conducted an unbiased survey of a selection of blocks using alternative mutagenesis schemes. For simplicity, these results are incorporated into the first section of the manuscript, even though machine learning models are only introduced later.

Machine learning predictions of gain-of-binding and inconsistent staining in blocks overlapping the same predicted binding motif led us to suspect that three transition block mutations caused a simultaneous loss and gain-of-binding. Using alternative GC-preserving transversion mutagenesis scheme (HT2 block 14) and targeted 2bp mutations (FL block 4, NEU3 block 8), we concluded that was likely the case and updated our block assessment accordingly (Supplementary Figure 2A).

An additional unbiased survey of 13 blocks (1 gain, 2 wild-type, 6 minor loss, 4 major loss) using alternative GC-content preserving transversion and scrambling block mutagenesis schemes revealed no major differences, overall validating original transition scheme as primarily causing loss-of-binding (Supplementary Figure 2B). Specifically, in 9/13 cases transition (the default scheme for this study) matched the transversion or scramble result perfectly. In 2 of 4 remaining cases, the difference was very minor and resulted in no change in score. In particular, for FL block 12 scrambling mutagenesis induced a weak gain of heart staining, which is likely explained by an accidental introduction of a GATA motif. The rest of the staining pattern was identical between transition and scrambling mutagenesis. For FL blocks 7 and 10 the differences between transition and transversion mutagenesis were more pronounced, first one changing the direction of effect (from minor loss to minor gain) and the other only changing the magnitude (major loss score 3 to score 2; Supplementary Figure 2B). We conservatively decided to use the transition result as final functional block annotation in all 13 cases.

Finally, classification of HT2 block 16 was updated from wild-type (single block transition mutagenesis result) to minor loss. This was based on the fact that 1bp mutation of a predicted double NFI site overlapping that block had a strong additional loss-of-function effect in combination with 1bp mutation targeting SRF site (see Supplementary Figure 5B).

In conclusion, we updated the classification of four blocks as follows: FL block 4 gain −> minor loss, NEU3 block 8 major loss score 3 to major loss score 1, HT2 block 14 major loss score 2 −> full loss (score 0) and HT2 block 16 wild-type −> minor loss.

### Supplementary Note 3: Rejected best-fit models for enhancers NEU1 and NEU2. Alternative NEU3 models

Two enhancers with brain activity in transgenic assay were not included in the analysis of machine learning models results due to low correlation of model predictions with *in vivo* results (NEU1) or lack of tissue-appropriate models (NEU2). This supplementary note contains additional analysis of these enhancers and their models.

Best model for **NEU1** enhancer was derived from midbrain ATAC-seq dataset at E11.5, with R^2^=0.29 (Supplementary Figure 6A, FDR > 0.01, not significant), far below the worst best-fit model included in the main analysis (HT3, R^2^=0.5). Log2 fold change predictions from this NEU1 model had a relatively narrow range, with most predictions being above −0.3, compared to best-fit models predictions below −1. This implied limited sensitivity of NEU1 models (Supplementary Figure 6A).

Mutations in different blocks of NEU1 affected different brain structures specifically (fore-, mid- and hindbrain), which could explain poor correlations based on whole-embryo assessment of *in vivo* mutational effect (e.g. mutations that abolished either fore- or hindbrain activity would both be classified as “major loss”). We examined fore-, mid- and hindbrain models, looking for tissue-specific prediction outliers and did not find any that would explain the poor fit (Supplementary Figure 6A). The strongest prediction outliers were shared by all three models and involved gain-of-function mutation of block 21 (singly or together with other blocks), which did not have tissue-specific impact. We conclude tissue-specificity was not the main driver of poor model fit.

For completion, we examined the contribution score predictions of the midbrain NEU1 model. The prediction contained many isolated CG-dinucleotides that did not appear to be TF binding motifs, along with LHX8, E-box, SOX and PITX/OTX2 motif predictions (Supplementary Figure 6B). While LHX8 and E-box sites overlapped a major loss block, the SOX and PITX/OTX2 sites were found within wild-type blocks, calling these limited predictions into question. We speculate NEU1’s cell type(s) of activity is poorly represented in whole-tissue samples on which we have built our models, leading to poor correlations and limited predictive power.

All five significantly correlating models of brain-active **NEU2** enhancer were derived from liver and heart tissues (R^2^=0.51-0.57), in which this enhancer had no activity. The best neuronal/brain model for this enhancer was based on interneuron 4 cluster of brain scATAC dataset, with (statistically insignificant) R^2^ of just 0.1. Models based on bulk ATAC-seq and DHS brain datasets showed even lower correlations.

It could be speculated that NEU2 is active in a non-neuronal cell type that is rare in bulk brain samples, thus neither contributing enough signal to these samples, nor sharing the regulatory logic with the most common cell types in the brain. Further, it could be speculated that the cell type in which NEU2 is active shares some similarities with liver and heart samples. Therefore, it should be theoretically possible to learn aspects of NEU2’s function from its best-fit open chromatin liver model (R^2^=0.57; Supplementary Figure 6A).

Best-fit model predicted two isolated GATA sites and one GATA1-TAL site, accounting for 5/7 major or full loss blocks (Supplementary Figure 6B). Targeted 1bp mutation of TAL1 part of GATA1-TAL site yielded a similar result to the block mutation 17 that encompassed the TAL1 part (Supplementary Figure 6C, left), supporting the model. However, of the two block mutations overlapping the first isolated GATA site (19-20), only the first one resulted in major loss of function, while the other did not affect the expression of the construct (despite strong prediction of −0.78 log-fold change in signal). Considering the possibility that the results of mutagenizing the second block were confounded by a simultaneous loss of GATA and gain of another (hypothetical) site, we ablated the putative GATA site directly through 1bp mutation. This perturbation resulted in no change of activity (Supplementary Figure 6C, right), strongly arguing against GATA TF binding this site. With virtually all other predicted sites being GATA, this result called the entire model prediction into question. Finally, we also observed that the model made three more strong log2 fold change predictions of block effect (blocks 4, 13 and 18, absolute effects of 0.39 or more), which were not borne out by experimental data, further invalidating the model.

An independent TF motif scan suggested that two adjacent blocks 6-7, which did not have any contribution score predictions in the liver model, could bind a weak PITX/OTX2 site, in line with brain activity of the enhancer. An *in silico* saturation mutagenesis on the e11.5 midbrain ATAC-seq model supported that hypothesis, with PITX/OTX2-like contribution scores appearing upon *in silico* mutation that would strengthen the existing site (Supplementary Figure 6D, A68T). No other sites were discovered in that screen. Experimental introduction of a 1bp mutation designed to destroy the PITX/OTX2 site (A70G) largely recapitulated the effect of block mutations overlapping the PITX/OTX2 site. We conclude that NEU2 is unlikely to share a liver/heart GATA-driven logic, but may use a neuron-like logic, for which we currently lack a suitable model.

Interestingly, the third enhancer active in the brain in our study, **NEU3**, had open chromatin signal in neural tissues, but also in the face and limbs, in which no *in vivo* activity was observed. In other words, NEU3 appeared to be poised in face and limbs and to share some of their functional logic. Models derived from brain tissues made remarkably similar predictions to models derived from face and limb tissues. For example, correlation (R^2^) between contribution scores of the best-fit neural NEU3 model to contribution scores from limb and face datasets was 0.57-0.74, compared to correlation with other brain/neural datasets of 0.58-0.88. Conversely, “face E11.5 (ATAC)” (model with highest correlation of 0.8 to *in vivo* results for NEU3 out of face/limb models), had R^2^ of 0.54-0.92 to brain/neuronal datasets. This implies the same factors mediate chromatin openness in both tissue types, and that some, yet unidentified factor makes the enhancer active only in the brain (or specifically inactive in the face and limbs).

Altogether, these results indicate that transcriptional activity can sometimes be learned from open chromatin signal at poised loci, but caution and experimental confirmation is needed in such cases. Until tissue-appropriate, activity-based models are trained, this form of “transfer learning” may be practically useful for prioritizing experiments and fine mapping of human variants.

### Supplementary Note 4: Systematic assessment of signal change based machine learning model predictions

We used machine learning models primarily to find TF binding motifs, which are revealed by the contributions scores. The models were evaluated on their ability to detect a site in reference sequences consistent with functional annotation of the block. However, the models could theoretically correctly predict the effect of introduced mutations without detecting the presence of a binding motif. Furthermore, in our model assessment we did not take into account the creation of novel binding motifs, which would only be present in the contribution score tracks of the mutated, but not the reference sequence. This Supplementary plementary Note provides an additional analysis of the five enhancers and their best-fit models, using direct model predictions of signal change.

We predicted signal change for each single block transition mutation. Since these predictions are continuous, we binarized them using a threshold. We picked an absolute log2 fold change signal cutoff of 0.32, corresponding to 25% change, so as to correctly classify at least 90% of wild-type blocks. For FL, we used the most extreme (absolute) prediction of the three selected models (limb, hindbrain and glutamatergic neurons). We referred to this prediction set as the “cutoff method” and compared it to the “contribution method”, the final set of reference sequence binding motif predictions, which includes alternative models and degenerate binding motifs.

Overall, the cutoff method performed similarly to the contribution method, with higher fraction of correctly classified major loss blocks (78% vs 69%), but lower of minor loss (31% vs 38%) and gain blocks (20% vs 60%; Supplementary Table 5), while maintaining a similar specificity (92% vs 94% of correctly classified wild-type blocks).

**Supplementary Table 5.**
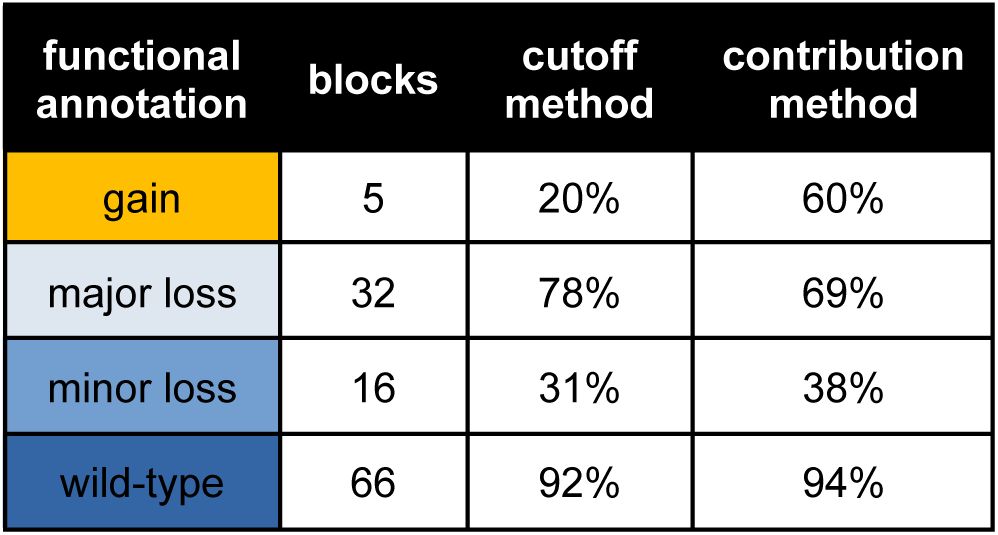
Machine learning model method comparison. Percentages are fraction of the blocks with a given functional annotation that were correctly classified by each of the methods.

Discrepancies between these two methods involved 16 blocks, primarily in enhancers FL and HT1 (6 blocks each). The majority of discrepancies (9/16) involved cutoff method predicting a change, where contribution method predicted no binding motif. In 7 out of these 9 cases, the cutoff method was correct. This implies models contained some information that could not be extracted using contribution scores. Similar “hidden information” was extracted by us as degenerate TF motifs using saturation mutagenesis (see Figure 3), but the result above implies more remains to be discovered.

The remaining 7 cases involved two blocks with correct contribution prediction (MEIS-TEAD degenerate site in HT1), one with incorrect prediction (degenerate MEF2 site in HT2) and four more complex cases (FL blocks 4-6 and HT2 block 14). One complex example involved mutagenesis of FL block 4. This block mutation likely resulted in a simultaneous creation of a TWIST1/HAND2 activator motif and destruction of a SOX activator motif (see Supplementary Figure 2A). With overall outcome being a gain of function, we speculate that the novel TWIST/HAND2 site contributed more to enhancer activity than was lost by ablation of the SOX site. Targeted destruction of the SOX motif through 2bp mutagenesis confirmed that the “true” functional annotation of this block is minor loss of function. While the cutoff method was technically correct in predicting the outcome of block mutagenesis, the contribution method predicted the functional annotation. Contribution method prediction was correct by accident, since for this method we only considered the presence of an activator SOX motif, but not the gain of TWIST1/HAND2. In another complex case, mutagenesis of HT2 block 14 resulted in destruction of a predicted SRF motif and creation of a novel SP/KFL site. The phenotypic result was major loss of function, implying that SRF contributed more to HT2 activity than the novel SP/KFL motif could compensate for, the opposite of FL block 4 case. The cutoff method incorrectly predicted this mutation will lead to no change of function. Again, the contribution method was correct here by accident, as gain of a novel SP/KFL motif was not taken into account when using this method. If it was, the result would be ambiguous, as one activator site being replaced by another one cannot be easily interpreted in terms of overall direction of change, without making assumptions about relative magnitude of contribution scores.

We conclude that predictions of mutation effects based on signal change (“cutoff method”) overall yielded results more closely aligned with outcomes of our experiments than predictions of binding motifs in reference sequences (“contribution method”). This was for the most part due to contribution scores not detecting binding motifs where the model strongly and correctly predicted a change of function. In practice, both methods complement each other, since signal change needs to be interpreted as either gain or loss of binding by the contribution scores and contribution scores may sometimes be unable to extract information available to the model.

## Methods

### Transgenic assay

Transgenic E11.5 mouse embryos were generated as described previously^9^. Briefly, super-ovulating female FVB mice were mated with FVB males and fertilized embryos were collected from the oviducts. Enhancer sequences were synthesized by Twist Biosciences and cloned into the donor plasmid containing minimal Shh promoter, lacZ reporter gene and H11 locus homology arms (Addgene, 139098) using NEBuilder HiFi DNA Assembly Mix (NEB, E2621). The sequence identity of donor plasmids was verified using long-read sequencing (Primordium). Plasmids are available upon request. A mixture of Cas9 protein (Alt-R SpCas9 Nuclease V3, IDT, Cat#1081058, final concentration 20 ng/μL), hybridized sgRNA against H11 locus (Alt-R CRISPR-Cas9 tracrRNA, IDT, cat#1072532 and Alt-R CRISPR-Cas9 locus targeting crRNA, gctgatggaacaggtaacaa, total final concentration 50 ng/μL) and donor plasmid (12.5 ng/μL) was injected into the pronucleus of donor FVB embryos. The efficiency of targeting and the gRNA selection process is described in detail in Osterwalder 2022^9^.

Embryos were cultured in M16 with amino acids at 37°C, 5% CO_2_ for 2 hours and implanted into pseudopregnant CD-1 mice. Embryos were collected at E11.5 for lacZ staining as described previously^9^. Briefly, embryos were dissected from the uterine horns, washed in cold PBS, fixed in 4% PFA for 30 min and washed three times in embryo wash buffer (2 mM MgCl2, 0.02% NP-40 and 0.01% deoxycholate in PBS at pH 7.3). They were subsequently stained overnight at room temperature in X-gal stain (4 mM potassium ferricyanide, 4 mM potassium ferrocyanide, 1 mg/mL X-gal and 20 mM Tris pH 7.5 in embryo wash buffer). PCR using genomic DNA extracted from embryonic sacs digested with DirectPCR Lysis Reagent (Viagen, 301-C) containing Proteinase K (final concentration 6 U/mL) was used to confirm integration at the H11 locus and test for presence of tandem insertions^9^. Only embryos with donor plasmid insertion at H11 were used. The stained transgenic embryos were washed three times in PBS and imaged from both sides using a Leica MZ16 microscope and Leica DFC420 digital camera.

### Correlating predictions of machine learning models and *in vivo* results

To assess fit between the models and *in vivo* results, experimental results were scored on a scale from 0 to 7, with 0 indicating full loss of function, 1-4 indicating various degrees of major loss of function, 5 indicating minor loss of function, 6 indicating no change, 7 a gain of function. The Spearman correlation (R) between this *in vivo* score and model predicted log2 fold change in open chromatin signal across all single and multi-block transition mutagenesis constructs was computed across for each model and enhancer combination. Total predicted signal for the input sequence was used. All model estimates were obtained from the count head, using as input 2114 bp centered on the enhancer, flanked by the reporter construct (H11 locus left homology arm on the left and Shh promoter and reporter LacZ gene on the right).

### Sensitivity, specificity and estimation of binding site numbers

Sensitivity and specificity were calculated simply as fractions of, respectively, functional or wild-type blocks overlapping model-predicted motifs. Positive predictive value was calculated as a fraction of predicted binding motifs overlapping at least one functional block. The GATA motif in HT1 block 5 (classified as major loss) also overlapped block 6 (gain) by 1bp, which was ignored for the sake of simplicity.

To obtain the model-corrected number of binding sites per enhancer, we counted each predicted binding site once (even if it spanned multiple blocks) and assumed that each functional block without a prediction contains exactly one site - an activator one, if loss of function was observed upon block mutagenesis or a repressor, if gain of function was observed. We excluded sites predicted to be in non-functional blocks.

### Selection of paired block mutations

We selected only block pairs that were separated by at least 1 block, to avoid potential gain-of-binding events at the interface of mutagenized blocks. We also excluded combinations of full loss of function blocks with other loss of function blocks, since the likely outcome - full loss of function - cannot be classified as either additive or non-additive in a meaningful way.

### Machine learning models training and interpretation

Training of scATAC ChromBPNet models included in this study was described previously^19,43^.

Reference genome (mm10), blacklist regions, filtered BAM files for pair-end data and unfiltered BAM files for single-end data (ATAC-seq and DHS) were obtained from the ENCODE portal^31,32^. For unfiltered BAM files, an additional filtering step was performed using ‘samtools view -b -@50 -F780 -q30’. Isogenic replicates for each biological sample were merged to yield consolidated reads. For ATAC-seq samples, the peaks were directly retrieved from the ENCODE portal. For DHS samples, we used MACS2^44^ and followed the ENCODE ATAC-seq protocol for peak-calling. We further removed regions that overlap with blacklist regions. The dataset was divided into three groups (training, validation, and testing) by chromosome (1-19, X and Y). We employed a 5-fold chromosome hold-out cross-validation approach with different sets of chromosomes for different groups in each fold. Group compositions for each fold are available <u>here</u>.

ChromBPNet models^19^ were trained to predict the read counts given 2114 bp sequences from both peak and background regions, or from background regions alone. The ultimate output of ChromBPNet was a prediction of counts corrected using background region model for Tn5 enzyme effects. The background regions were chosen not to overlap peak regions, to have fewer reads than a minimum number of total counts observed in any peak region and to match the GC-content distribution of peak regions. Pearson correlation between predicted and observed log counts was used as a metric of fit during training. We utilized the DeepSHAP implementation of the DeepLIFT algorithm to derive base-resolution contribution scores for each input sequence^20,45^.

Motifs were identified using web server TOMTOM version 5.5.6 with default settings^46^.

## Acknowledgements

This work was supported by a U.S. National Institutes of Health (NIH) grant to L.A.P. (R01HG003988). Research was conducted at the E.O. Lawrence Berkeley National Laboratory and performed under U.S. Department of Energy Contract DE-AC02-05CH11231, University of California (UC). The authors acknowledge funding support from NIH grants 5U24HG007234, U01HG009431, and U01HG012069 to A.K. We would like to thank Laura Cook, Evgeny Kvon, Om Patange and Fabrice Darbellay for critical reading of the manuscript.

## Contributions

Mi.K. designed the study, analyzed the data and wrote the manuscript. B.Z., A.P. prepared the machine learning models and ran the predictions. I.P.-F., C.S.N., S.T. and Mo.K. performed microinjections and surgical embryo transfers. R.D.H., K.v.M., S.B., E.B. and Y.Z. prepared the constructs and genotyped the embryos. J.A.A., genotyped and imaged the embryos, and supervised the technical team. A.K. supervised B.Z. and A.P and obtained funding for running of the machine learning predictions. D.E.D. designed the study and provided general supervision. A.V. designed the study, provided general supervision, obtained funding and contributed substantially to writing of the manuscript. L.A.P. designed the study, provided general supervision and obtained funding.

